# Pulsed-SILAC in single mouse embryos reveals early embryonic protein synthesis dynamics and phosphosite regulation

**DOI:** 10.64898/2026.09.24.754052

**Authors:** Luisa Marie Schmidt, Maico Y. Lechner, Javier Martin-Gonzalez, Jason A. Halliwell, Ivan Mikicic, Molly Pleasants Lowndes, Ivo Alexander Hendriks, Katrin Stuber, Mads Lerdrup, Eva Hoffmann, Jesper V. Olsen

## Abstract

Early embryogenesis relies extensively on maternally deposited products until zygotic genome activation, yet the dynamics for the synthesis of new proteins in mammalian embryos remains poorly characterized. To address this, we applied pulsed stable isotope labelling by amino acids in cell culture (pSILAC) combined with narrow-window data-independent acquisition mass spectrometry to single mouse oocytes and embryos to resolve *de novo* protein synthesis during early embryogenesis. This revealed that the maternal proteome is not a static reservoir, with components of the subcortical maternal complex and amino acid transporters SLC7A1/2 being actively synthesized during the earliest developmental stages. Furthermore, phosphoproteomic analysis identified hundreds of previously unreported phosphosites and extensive regulation during the oocyte-to-embryo transition. Notably, phosphorylation of the PRC2-interacting KLP motif of EZHIP emerged as a potential regulatory mechanism, with modification of this region reducing EZHIP-PRC2 interaction and coinciding with H3K27me3 remodelling. Together, single embryo pSILAC revealed a maternal proteome that is continuously synthesized, recycled, and post-translationally regulated during early embryogenesis.

## Introduction

Early embryogenesis is a critical period marked by tightly regulated cellular divisions, orchestrated developmental transitions, and profound changes in gene expression.[1–3] In mammals, this phase has traditionally been considered largely dependent on maternally deposited mRNAs, proteins, and metabolites, which drive early developmental processes until the zygotic genome becomes transcriptionally active.[4, 5] In the mouse, zygotic genome activation (ZGA) occurs in two waves, minor activation during the late 1-cell stage and a more robust major activation at the 2-cell stage.[2, 6] While the presence of maternal and zygotic mRNAs has been extensively profiled using transcriptomic approaches in mouse and human cells [7–9], it is evident that transcript abundance does not necessarily correlate with protein expression due to translational regulation, post-transcriptional modifications, and selective mRNA decay [10, 11]. Specific waves of protein synthesis are required for processes such as cell cycle progression, chromatin remodelling, epigenetic reprogramming, and the establishment of pluripotency.[12–14] However, our understanding of protein synthesis dynamics during early development remains limited, largely due to technical challenges. Traditional proteomics approaches often lack the sensitivity to detect low-abundance proteins in the scarce material available from early embryos and cannot differentiate between newly synthesized proteins and their turnover rates.

To overcome these limitations, metabolic labelling strategies have emerged as powerful tools to study protein dynamics. Among these, pulse-chase Stable Isotope Labelling by Amino acids in Cell culture (pSILAC) provides a sensitive and direct method to measure newly synthesized proteins.[15–18] By culturing cells in media with heavy isotope-labelled amino acids for defined periods, pSILAC enables incorporation of labelled amino acids into newly synthesized proteins.[19] These can then be distinguished from pre-existing proteins using high-resolution mass spectrometry, allowing precise quantification of protein synthesis over time.

While pSILAC has been extensively used in cultured cells and some model organisms, its application to mammalian embryos has been limited. Recent advances in miniaturized proteomic workflows, such as optimized sample preparation protocols [20–22], increased sensitivity of mass spectrometers [23–25], and software development[26, 27], now make it feasible to apply pSILAC to single cells from human cell lines [28] as well as early-stage mouse embryos. This opens new avenues for investigating the temporal regulation of protein synthesis during preimplantation development, and for identifying protein networks that are dynamically regulated before and during ZGA.

In this study, we applied a pSILAC-based proteomic strategy to early mouse embryos to capture the synthesis of new proteins across key developmental windows. By switching embryos between light and heavy SILAC media at defined time points and analysing them using data-independent acquisition (DIA) mass spectrometry, we provide a high-resolution view of protein synthesis dynamics during the transition from the zygote to the morula stage. This approach allows us to identify newly synthesized proteins, characterize stage-specific synthesis patterns, and uncover regulatory mechanisms that drive early embryonic development in mice and potentially in humans.

## Results

### Pulsed SILAC resolves protein synthesis dynamics during early embryogenesis

To investigate protein turnover dynamics during the early transition from zygote to morula, fertilized mouse eggs were isolated and cultured in either light (L) or heavy (H) Stable Isotope Labelling by Amino Acids in Cell Culture (SILAC) medium. The light medium contained [¹²C₆, ¹⁴N₂]-L-lysine and [¹²C₆, ¹⁴N₄]-L-arginine, while the heavy medium contained [¹³C₆, ¹⁵N₂]-L-lysine and [¹³C₆, ¹⁵N₄]-L-arginine.[18] After 24 h of culture, embryos reached the 2-cell stage, at which point the medium were switched to either light or heavy SILAC medium.

This resulted in four labelling conditions: H◊H, H◊L, L◊H, and L→L, each represented by approximately 20 replicates (**Figure 1A**). No developmental differences were observed between control embryos cultured in standard KSOM medium supplemented with BSA (**Figure 1B**) and SILAC-labelled embryos (**Figure 1C**). Morulae proteomes were extracted and digested using the One-Tip protocol and analysed by liquid chromatography-tandem mass spectrometry (LC-MS/MS) using the Evosep One LC coupled to the Orbitrap Astral mass spectrometer operated in narrow-window data-independent acquisition (nDIA) mode (**Figure 1D**).[21, 24]

**Figure 1.**
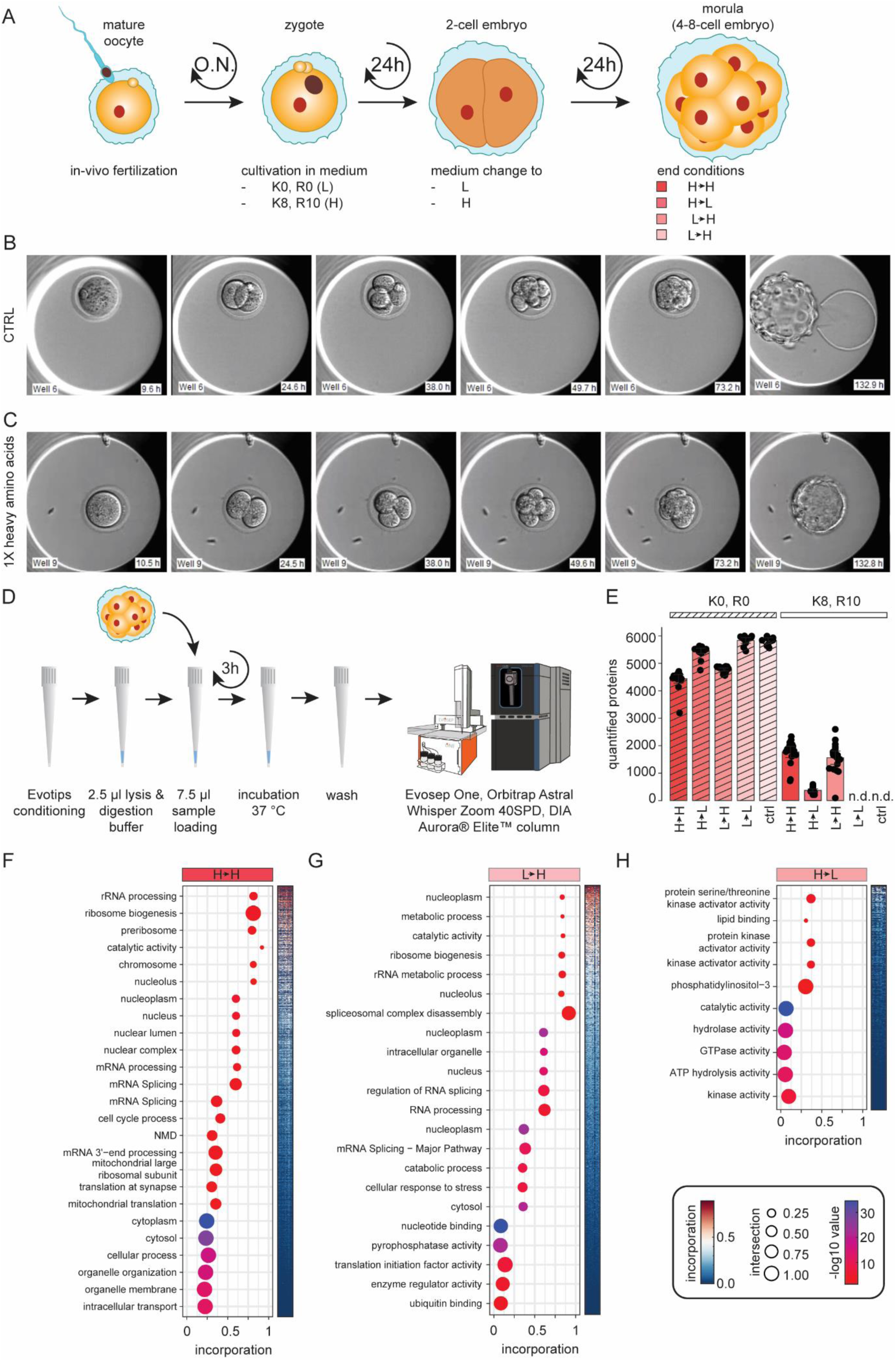
Single embryos show incorporation within the first 48 h. **A**) Schematic overview of the workflow for the cell culture experiment. From left to right: zygotes were isolated after natural mating and cultured in either light [K0, R0] or heavy [K8, R10] SILAC medium. After 24 h of incubation, the medium was switched (to light or heavy), and embryos were harvested after a total of 48 h, at the morula stage. **B**, **C**) Representative EmbryoScope pictures from the development of the first 130 h of zygotes in control medium (**B**) and in medium containing heavy amino acids (**C**). **D**) Morulae were loaded onto preconditioned Evotips and spun down into the 3.5 µl of lysis and digestion buffer. After washing, samples were analysed by LC-MS/MS (**B**). **E**) Number of protein groups identified in the light and heavy SILAC channels using DIA-NN for data analysis. **F**-**H**) Top identified Gene Ontology terms for the incorporation, separated to quartiles and for the three different conditions of H ◊ H (**F**), L ◊ H (**G**), and H ◊ L (**H**). Significant proteins were selected by g:SCS algorithm a=0.05. CTRL, control; DIA, data independent acquisition; H, heavy; LC-MS/MS, liquid chromatography tandem mass spectrometry; L, light; SILAC, stable isotope labelling in cell culture; O.N., overnight; SPD, samples per day.

Prior to the main experiment, digestion conditions for the One-Tip protocol were optimized using HeLa cells in the range of 250-4000 cells, approximating the size of oocytes to 8-cell-stage embryos. Two protein extraction and digestion (master mix) formulations were tested, with master mix 1 containing four times more Lys-C and trypsin proteases than master mix 2. Both were evaluated at digestion times of 15 minutes and 3 h. Master mix 1 outperformed mix 2 under short digestion conditions; however, no significant differences were observed after 3 h of digestion, despite a higher rate of missed cleavages in master mix 1 (**SI Figure 1A-D**). Similar trends were observed in data-dependent acquisition (DDA) mode, although only one-fifth of the peptides identified using DIA were also detected in DDA (**SI Figure 1E-H**). To better compare DIA and DDA performance in a pulsed-chase SILAC (pSILAC) context, HeLa cells were cultured in light or heavy SILAC medium, and the medium was switched after 4 h to mimic the embryo conditions. A total of 2000 cells (approximately equivalent to a 4-cell-stage embryo) were processed using the One-Tip protocol for either 15 min or 3 h. Longer digestion times significantly improved the identification of heavy-labelled peptides (**SI Figure 1I-P**). Based on these findings, master mix 1 with a 3 h digestion time and DIA-based analysis was used for all subsequent experiments.

In morulae, between 4500 and 6000 proteins were quantified in the light-SILAC channel, and 1000-2000 proteins were quantified in the heavy-SILAC channel using DIA-NN software for raw LC-MS data analysis. As expected, no heavy-SILAC signal was detected in the L→L condition or in the KSOM only control (**Figure 1E** and **SI Figure 2A, B**). Calculation of heavy amino acid incorporation rates for newly synthesized proteins revealed average incorporations of 25%, 9%, and 26% for the conditions H◊H, H◊L, L◊H, respectively (**SI Figure 2C**). Comparison of the three conditions revealed approximately 600 newly synthesized proteins common to all conditions, while H◊H and L◊H shared 1687 newly synthesized proteins, suggesting that protein synthesis increases substantially after the 2-cell stage (**SI Figure 2D, E**). Supporting this hypothesis, gene ontology (GO) term enrichment analysis of proteins with the highest incorporation for H◊H and L◊H showed overrepresentation of ribosome biogenesis and RNA metabolism (**Figure 1F, G**).[29, 30] Additionally, proteins localized to the nucleoplasm and chromosome showed high heavy-SILAC incorporation, while subcellular compartment analysis indicated that the nuclear envelope had the highest H/L ratio, and the proteasome displayed the most proteins with a negative log2 ratio across all conditions (**SI Figure 2F, G**). Interestingly, during the first 24 h, proteins with the highest incorporation were predominantly associated with kinase activity, suggesting that these key cell signalling proteins are rapidly synthesized to initiate or regulate the onset of protein synthesis. (**Figure 1H**). While these results confirm that the setup can successfully identify newly synthesized proteins, the overall proportion of newly synthesized proteins was relatively low, most likely due to dilution of heavy amino acids in the zygote and the limited amino acid uptake during early developmental stages.

**Figure 2.**
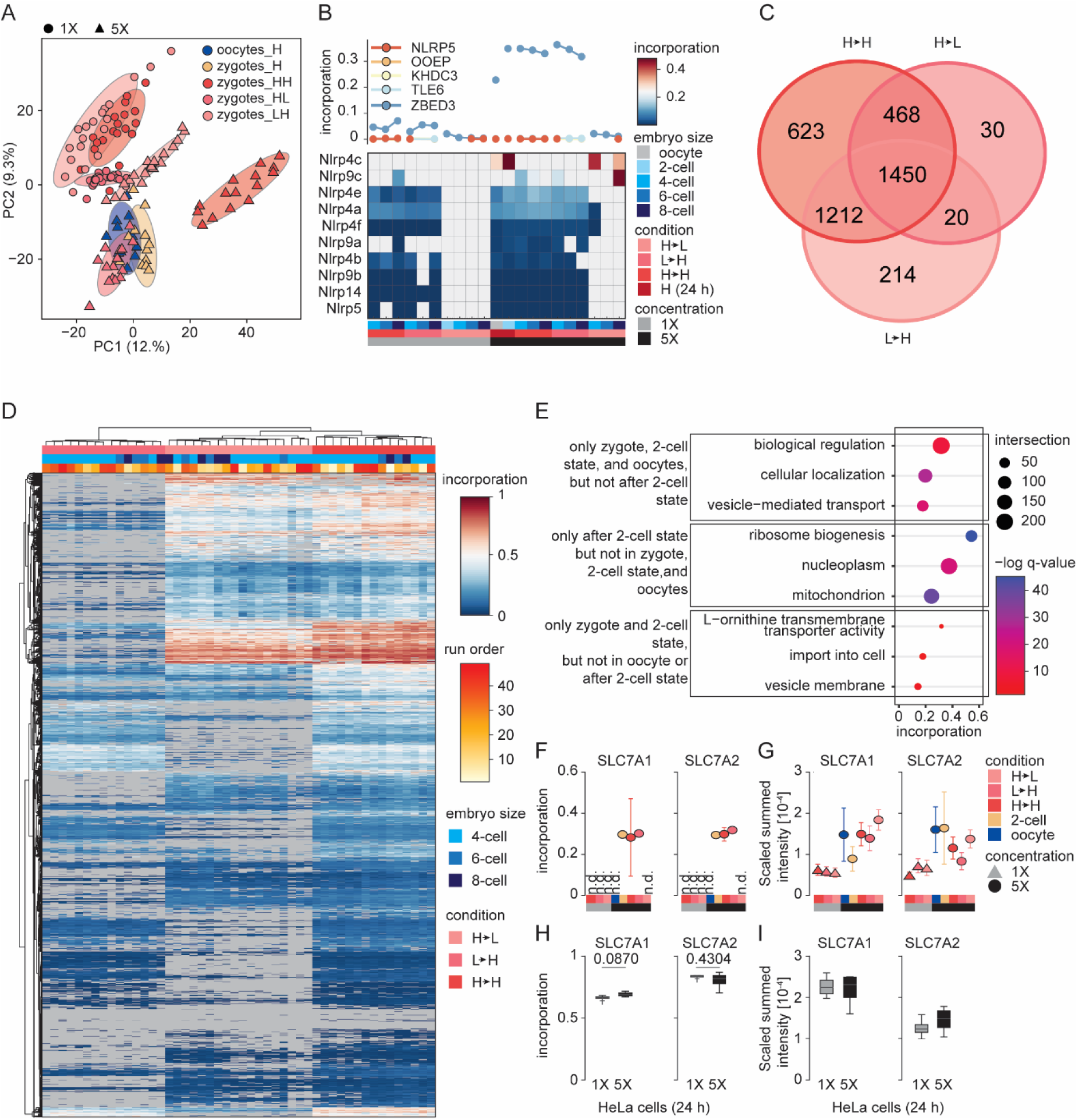
Increased heavy arginine and lysine concentrations enhance heavy-label incorporation and reveal changes in cationic amino acid transporters. **A**) Principal component analysis (PCA) of incorporation levels in oocytes (24 h), 2-cell embryos (24 h), and embryos (48 h) cultured with 1X and 5X concentrations of heavy amino acids under different labelling conditions (H◊H, L◊H, and H◊L) shows clear separation between the 5X conditions, but less distinction among the 1X conditions. **B**) Line plot showing incorporation in the oocyte complex for 1X and 5X amino acid concentrations and heatmap of NLRP proteins. Both indicate active protein synthesis during the first 24 h, but not during the second 24 h. **C, D**) Venn diagram (**C**) and heatmap (**D**) of newly synthesized proteins identified in the H◊H, H◊L, and L◊H conditions under the 5X concentration. **E**) Dot plot showing selected GO terms enriched in proteins synthesized during the first 24 h in fertilized and unfertilized eggs, only during the second 24 h, or exclusively in fertilized eggs during the first 24 h. Significant GO terms were selected by g:SCS algorithm a=0.05. **F, G**) Incorporation (**F**) and scaled summed intensity (**G**) of the high-affinity cationic amino acid transporter SLC7A1 and the cationic amino acid transporter SLC7A2, showing heavy amino acid incorporation restricted to the first 24 h in fertilized eggs in 5X concentration. Data is presented in mean ±SEM. **H, I**) Incorporation (**H**) and scaled summed intensity (**I**) of SLC7A1 and SLC7A2, showing no difference in incorporation between 1X and 5X within 24 h in HeLa cells. Data is presented in mean ±SEM. T-test doesn’t show any significance. PCA, principal component analysis; NLRP, NACHT, LRR, and PYD domains-containing protein; SLC7A1, high-affinity cationic amino acid transporter 1; SLC7A2, cationic amino acid transporter 2.

### Increased arginine and lysine concentrations enhance labelling and cationic amino acid transporter synthesis

To reduce the dilution effect of heavy amino acids, oocytes and zygotes were next cultured in medium containing five-fold (5X) concentrations of heavy-labelled arginine and lysine. Embryos developed normally and showed no morphological abnormalities compared to those cultured in standard (1X) heavy amino acid medium, indicating that elevated concentrations do not impair developmental progression (**SI Figure 3A**). Oocytes (mature oocytes cultured) and 2-cell embryos (zygotes d0.5 cultured to 2-cell embryos d1.5) were harvested after 24 h (*n*=12), and embryos at 48 h were collected as in the previous experiment (*n*=*13*). To further improve detection sensitivity, samples were analysed using the Evosep Eno LC coupled to an Orbitrap Astral Zoom MS, with the same setup as before (**SI Figure 3B**).[31, 32] Quantitative proteomic analysis revealed similar numbers of identified peptides and proteins in the light SILAC channel compared to the 1X experiment; however, the number of proteins detected in the heavy SILAC channel increased from approximately 2000 to 3000 under 5X conditions (**SI Figure 3C, D**). Histograms of incorporation values confirmed higher labelling efficiency at increased amino acid concentrations, with average increases of +0.05, +0.06, and +0.02 for the H◊H, H◊L, and L→H conditions, respectively (**SI Figure 3E**).

**Figure 3.**
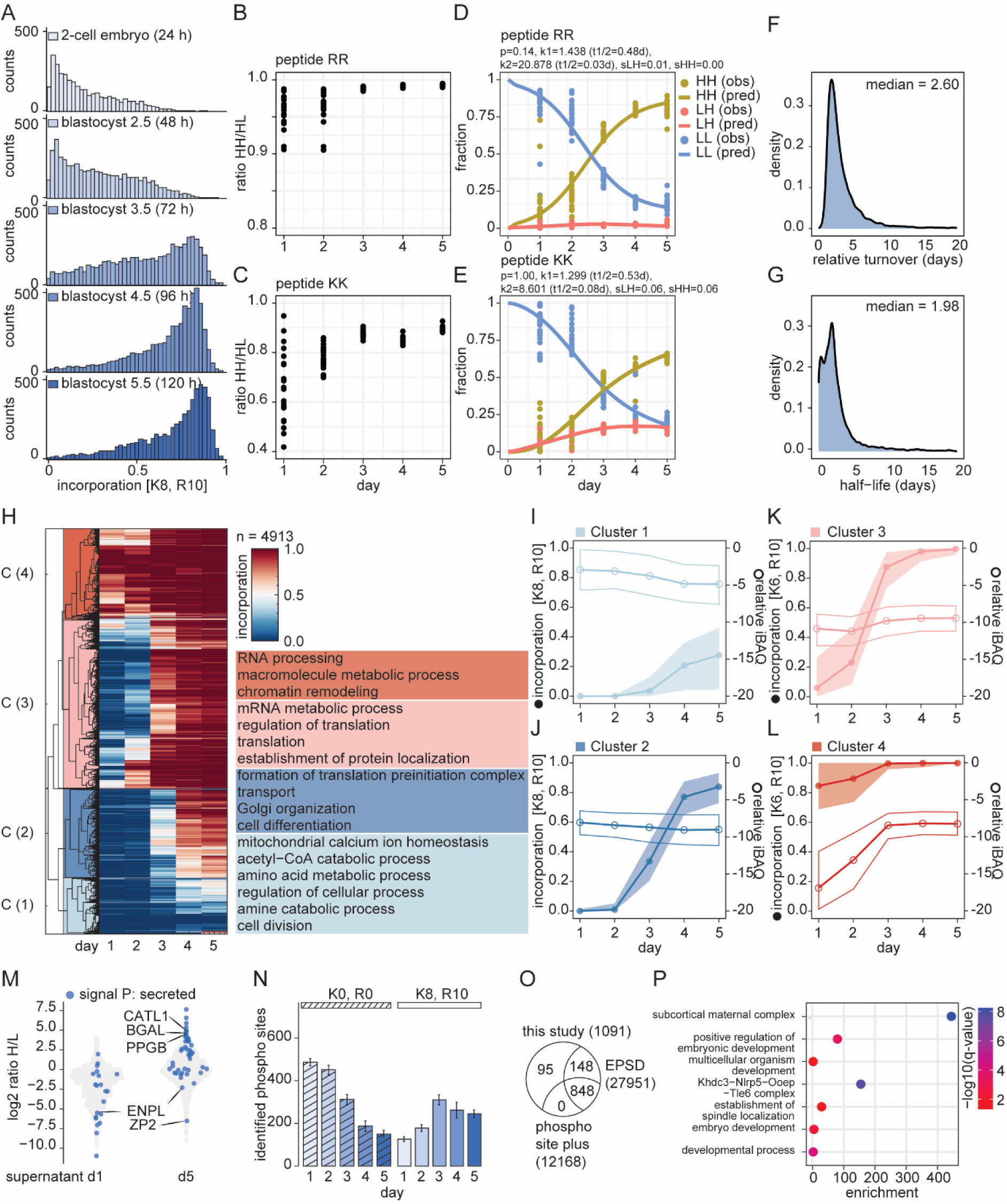
High recycling of lysine influences the calculated turn-over of proteins in embryos. **A**) Histogram of incorporation levels per counts across the five days shows an increasing incorporation of heavy amino acids with increasing time. **B, C**) Ratio of missed cleaved peptides containing two heavy (pepHH) and one light, one heavy (pepLH) arginine (**B**) or lysine (**C**). **D, E**) Fraction of pepHH (yellow), pepLH (red), pepLL (blue) compared to the total missed cleaved peptides containing either arginine (**D**) or lysine (**E**). **F, G**) Density curve for the half-life or proteins before correcting for recycling with a median of 2.6 (relative turnover) (**F**) and after correcting with a median of 1.98 (**G**). **H**) Hierarchical row clustering of mean corrected incorporation per day shows four specific clusters. Top selected GO terms are described on the right side of the heatmap. Significant GO terms were selected by Fisher’s exact test, Benjamini-Hochberg FDR correction. **I-L**) Incorporation and log2 relative iBAQ mean values of the four identified clusters. Outlines represent the standard deviation. On the left and right axis, incorporation and log2 riBAQ values are shown, respectively. **M**) Violin plot of the log2 H/L ration from the supernatant collected from the embryos at day 1-5. **M**) Identified phospho sites for light and heavy channels. **N**) Overlap of the identified phospho sites of the identified proteins with two phosphosite repositories (Phosphosite plus [43] and EPSD [44]). **O**) GO term enrichment analysis of the proteins carrying the newly identified 95 phospho sites. Significant GO terms were selected by g:SCS algorithm a=0.05. **P**) Density curve for the half-life or proteins using only phosphopeptides after correcting for recycling with a median of 2.69. **Q-T**) Incorporation of phosphopeptides compared to the protein. BGAL, beta-galactosidase gene; CATL1, carnitine palmitoyltransferase 1C; ENPL, Endoplasmin; EPSD, Eukaryotic Phosphorylation Site Database; H, heavy; iBAQ, intensity-Based Absolute Quantification; L, light; PPGB, Protective Protein for Beta-Galactosidase; ZP2, Zona Pellucida Glycoprotein 2.

Principal component analysis (PCA) of incorporation levels across embryos cultured under different labelling conditions (H◊H, L◊H, and H→L) demonstrated clear separation among the 5X samples, whereas the 1X samples showed less distinction between conditions (**Figure 2A**). Density scatter plots comparing incorporation between 1X and 5X concentrations across the same conditions revealed consistent trends and high Pearson correlations, confirming the reproducibility of the labelling strategy (**SI Figure 3F-H**). Correspondingly, bar plots indicated a 2-2.5-fold higher overall incorporation under 5X heavy amino acid conditions compared to 1X (**SI Figure 3I**). Thus, although the relative correlation to 1X conditions remained stable, the overall labelling efficiency was markedly improved. Closer examination of proteins associated with the subcortical maternal complex (SCMC) and the Nod-like receptor protein (NLRP) family revealed similar temporal patterns in both 1X and 5X conditions, with incorporation occurring primarily during the first 24 h and limited synthesis thereafter (**Figure 2B**). The SCMC multi-protein structure is essential for zygotes to progress beyond the first embryonic cell divisions as it helps manage processes like spindle placement, translation regulation, and epigenetic reprogramming.[33] While it is widely accepted that the SCMC is maternally deposited, we observed evidence of synthesis of the SCMC components during the first 24 h, implying possible translation of maternal mRNAs or limited zygotic contribution.[33–36] At the 5X concentration, Venn diagram and heatmap analyses of newly synthesized proteins across the H◊H, H◊L, and L→H conditions showed substantially greater overlap than in the 1X condition, with approximately 1500 newly synthesized proteins in common compared to only ∼600 in 1X (**Figure 2C** and **SI Figure 2D**). The heatmap revealed strong correlation among the three conditions with no association to LC-MS/MS run order (**Figure 2D**). Notably, clustering corresponded to embryonic cell stage, as 6-8-cell embryos consistently grouped together within each labelling condition. GO term enrichment analysis identified biological processes related to ribosome biogenesis and RNA metabolism during the second 24 h of development as overrepresented, whereas vesicle-mediated transport and cellular localization terms were shared between oocytes and embryos during the first 24 h (**Figure 2E**). Interestingly, during the first 24 h, only fertilized embryos and not oocytes showed enrichment for proteins associated with amino acid import and L-ornithine transmembrane transporter activity.

To investigate this observation in more detail, the high-affinity cationic amino acid transporter SLC7A1 and the cationic amino acid transporter SLC7A2 were examined across 1X and 5X conditions. SLC7A1 and SLC7A2 are the membrane proteins that specifically transport ornithine, arginine and lysine across cell membranes. Both transporters exhibited increased SILAC incorporation only under 5X conditions, indicating increased synthesis as a direct response to elevated arginine and lysine concentrations (**Figure 2F, G**). This finding is notable, as amino acid uptake in early embryogenesis is known to rely primarily on maternal transporters and become more active in the later blastocyst stage, which requires zygotic genome activation.[37]

In contrast, in HeLa cells cultured under equivalent 1X and 5X SILAC conditions, no significant increases in SLC7A1 or SLC7A2 incorporation were observed within 24 h (**Figure 2H, I**). This suggests that the response to increased amino acid availability is context-dependent and particularly pronounced in early embryos rather than representing a general consequence of increased medium concentrations.

### Lysine recycling influences protein turnover estimates during early embryogenesis

To further investigate protein turnover during early embryonic development, zygotes were cultured for up to five days in heavy medium, harvested, and analysed as described previously (**SI Figure 4A, B**). Heavy amino acid incorporation increased markedly after 96 h, which was also reflected in the number of identified proteins in the heavy SILAC channel and in the total protein identifications, averaging approximately 6500 and 7000 proteins per embryo in each SILAC channel, respectively (**Figure 3A** and **SI Figure 4B, C**).

**Figure 4.**
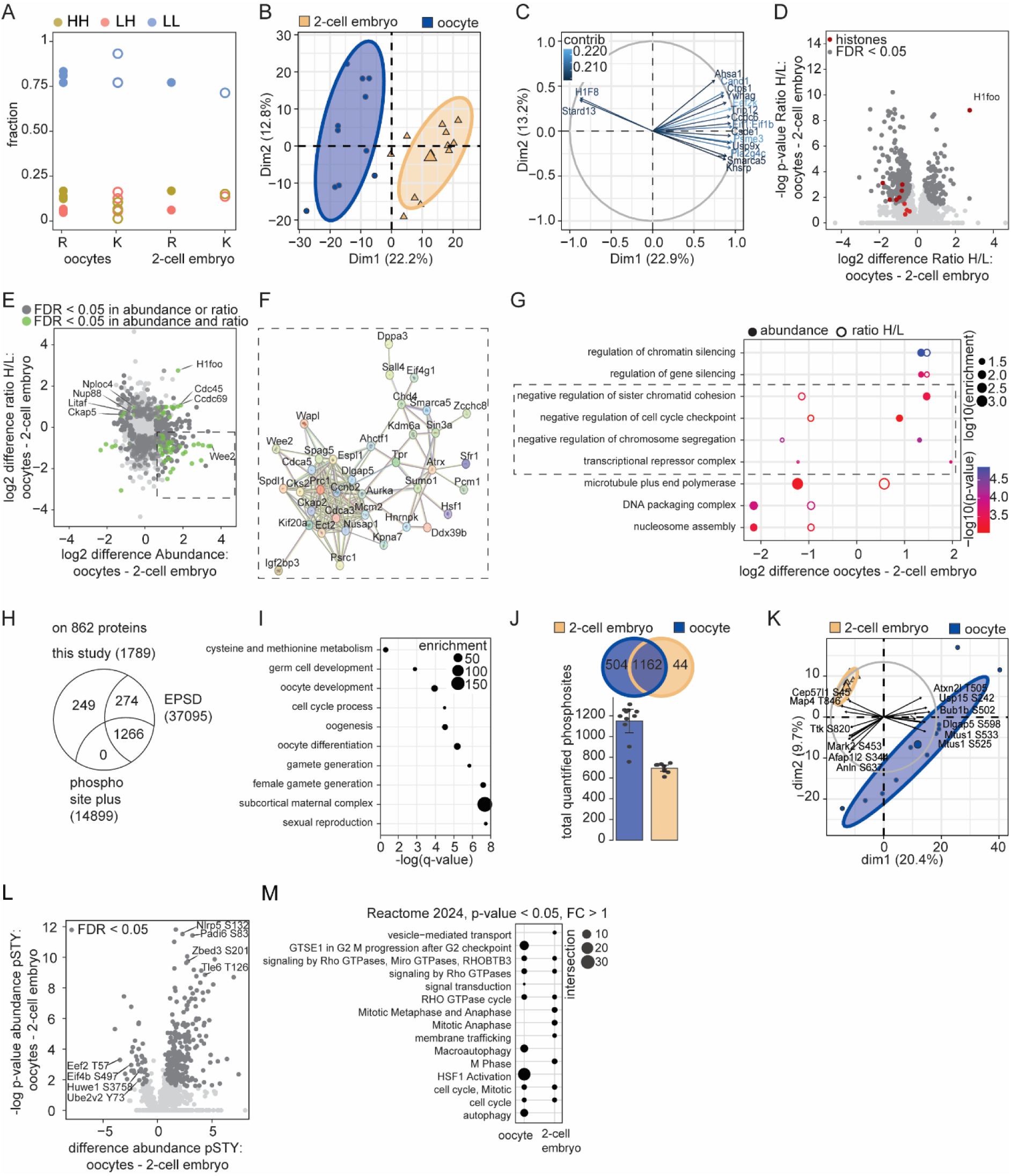
Oocytes and 2-cell embryos show distinct proteome and phosphoproteome profiles. **A**) Fraction of pepHH (yellow), pepLH (red), pepLL (blue) compared to the total missed cleaved peptides containing either arginine or lysine in oocytes and 2-cell embryos. **B, C**) PCA of projection (**B**) and loadings (**C**) on the protein abundances in oocytes (blue) and 2-cell embryos (orange). **D**) Volcano plot of the log2 differences in H/L ratios of oocytes and 2-cell embryos. Significant proteins (Permutation’s based FDR <0.05) and histones are marked in dark grey and red, respectively. **E**) Scatter plot of the log2 differences in abundance and H/L ratios of oocytes and 2-cell embryos. Significant proteins (Permutation’s based FDR <0.05) identified in only one, or both analyses are marked in dark grey and green, respectively. **F**) String network analysis of the defined proteins identified as significantly higher abundant with a lower ratio in oocytes compared to 2-cell embryos. **G**) Dot plot of the GO term analysis of the significantly abundant and newly synthesized proteins between oocytes and 2-cell embryos. **H**) Venn diagram of the identified phosphosites in this study compared to the two main phosphosite repositories phosphosite.org and EPSD on the identified 862 proteins.[43, 55] **I**) Dot plot of the GO term enrichment analysis of the proteins with the identified phosphosites only identified in this study. Significant GO terms were selected by g:SCS algorithm a=0.05 **J**) Bar plot of the number of identified phosphosites in oocytes (blue) and 2-cell embryos (orange). **K**) PCA of projection and loadings on the phosphosite abundances in oocytes (blue) and 2-cell embryos (orange). **L**) Volcano plot of the log2 differences in protein abundances in oocytes and 2-cell embryos. Significant proteins (Permutation’s based FDR <0.05) are marked in dark grey and interesting proteins and phosphosites are visualized by name. **M**) Dot plot of the Reactome analysis of significantly different abundant phosphosites between oocytes and 2-cell embryos. Significant GO terms were selected by g:SCS algorithm a=0.05. CCDC69, Coiled-Coil Domain Containing 69; CDC45, Cell Division Cycle 45; CKAP5, cytoskeleton-associated protein 5; EPSD, Eukaryotic Phosphorylation Site Database; FC, fold change; FDR, false-discovery rate; H1FOO/H1F8, H1 histone family, member O, oocyte-specific; L, light; LITAF, Lipopolysaccharide Induced TNF Factor; STARD13, StAR Related Lipid Transfer; NPLOC4, Nuclear protein localization protein 4 homolog; NUP88, nucleoporin 88; WEE2, Wee1-like protein kinase.

To correct for the recycling or reuse of light amino acids, derived from maternal deposits or protein degradation, missed-cleaved tryptic peptides were used as a proxy.[38–41] Ratios of peptides containing two heavy amino acid residues (pepHH) versus those containing one heavy and one light residue (pepHL) were determined for both lysine- and arginine-containing peptides. Notably, during the first two days, the pepHH/pepHL ratio was lower for both amino acids, indicating a higher degree of recycling of light residues (**Figure 3B, C**). This effect was particularly pronounced for lysine-containing peptides, which exhibited extensive recycling during the first 48 h. For arginine-containing peptides, pepHL species almost disappeared after day 3, likely reflecting arginine’s preferential use as a precursor for nitric oxide synthesis, whereas lysine is more frequently incorporated into ribosomal proteins.[42] To quantify this effect, amino acid-specific recycling rates were estimated and applied to the main dataset separately for lysine- and arginine-containing peptides (**Figure 3D, E**).[39] Because of the relatively high recycling rate observed for lysine in the early embryo cells compared to HeLa cells, the estimated median protein half-life decreased from 2.6 to approximately 2.0 days (**Figure 3F, G**). Both PCAs on the incorporation and protein abundance showed a clear separation between the early days and the late two days (**SI Figure 4 D, E**). Hierarchical clustering of mean incorporation per day revealed four major clusters, with Cluster 4 showing the highest protein turnover. GO term enrichment analysis identified Cluster 4 proteins as being involved in RNA processing and macromolecule metabolic processes (**Figure 3H**). In contrast, Clusters 1 and 2, which displayed lower incorporation rates over time, were enriched for proteins involved in mitochondrial homeostasis, acetyl-CoA catabolic processes and transport, and Golgi organization, respectively.

To determine whether changes in turnover reflected altered protein abundance or degradation, Intensity-Based Absolute Quantification (iBAQ) values were calculated and normalized to total protein intensity per sample (riBAQ) (**SI Figure 4G, H)**. iBAQ is a label-free MS method used to estimate the relative and quasi-absolute abundance of proteins within a single biological sample.[40] The fraction of proteins belonging to Cluster 1 decreased over time, whereas those in Cluster 4 increased in abundance. Proteins in Clusters 2 and 3 remained relatively constant (**Figure 3I-L**). These results suggest that proteins associated with RNA processing, macromolecule metabolism, and chromatin remodelling account for the majority of newly synthesized protein content. Interestingly, Cluster 2 was characterized by a late increase in protein synthesis despite relatively constant total protein abundance and was significantly enriched for GO terms related to transport and vesicle-mediated processes. We therefore investigated whether newly synthesized proteins were released from the embryos into the extracellular environment. Culture supernatants were collected at the indicated developmental stages, trypsin-digested as described above, and analysed by LC-MS/MS. While the total number of proteins identified in the supernatant remained relatively constant from day 1 to day 5 (approximately 2,200-2,600 proteins), the H/L ratio progressively increased, indicating an increasing contribution of newly synthesized proteins to the extracellular proteome (**Figure 3M**).

To identify proteins potentially released through the classical secretory pathway and reduce the contribution of proteins originating from cellular debris, identified protein sequences were analysed by SignalP for N-terminal signal peptides. Only 6% of the proteins were annotated as secreted proteins, but among these with an active synthesis at day 5 were cathepsin L (CTSL), protective protein/cathepsin A (CTSA), and β-galactosidase (GLB1). Interestingly, all three are lysosomal proteins, suggesting a link between the increased synthesis of vesicle-associated proteins and lysosomal/secretory trafficking at day 5.

Extracellular lysosomal proteases may contribute to remodelling of the peri-embryonic environment. Cathepsin L has previously been detected in the secretome of mammalian preimplantation embryos and has been associated with embryo quality.[45, 46]

To further explore the relationship between protein turnover and post-translational regulation, phosphorylation sites were analysed by searching the raw DIA data with phosphorylation as a variable modification. In each embryo sample, between 200 and 500 phosphosites were identified in the light and heavy channels, respectively (**Figure 3N**). While total protein identifications steadily decreased in the light channel and increased in the heavy channel, phosphopeptide identifications declined in the heavy channel after day 3. In total, 1091 unique phosphorylated peptides were identified, corresponding to approximately 650 phosphoproteins. Comparison with public phosphoproteome repositories revealed that these proteins contain around 28,000 annotated phosphorylation sites.[43, 44]Remarkably, 95 of the phosphosites identified in this study have not been reported previously (**Figure 3O**). These newly identified sites were mainly found on proteins associated with the SCMC, embryonic development, and developmental processes (**Figure 3P**).

Half-lives were also calculated from the phosphosite-specific SILAC incorporation profiles after correction for amino acid recycling, resulting in a median phosphosite-derived half-life of 2.69 days (**SI Figure 5B**). Overlap between half-lives calculated from protein abundance and their corresponding phosphosite-specific peptide abundances showed strong correlation, with exceptions for a subset of proteins (**SI Figure 5A-C**).

**Figure 5.**
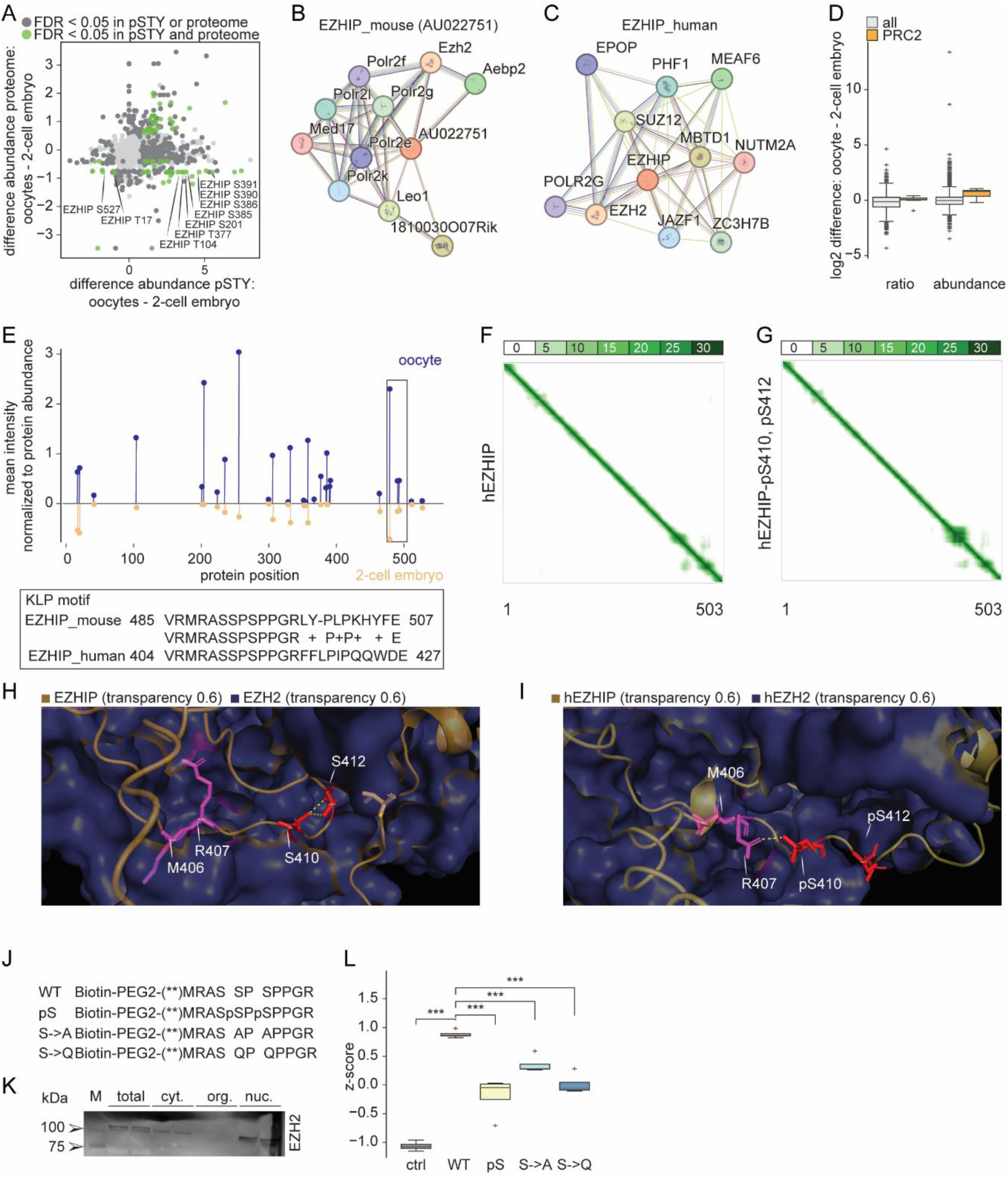
Phosphorylation within the EZHIP KLP region modulates its interaction with PRC2. **A**) Scatter plot of the log2 differences in abundance of the phosphosites and corresponding proteins in oocytes and 2-cell embryos. Significant proteins (Permutation’s based FDR <0.05) identified in only one, or both analyses are marked in dark grey and green, respectively. **B, C**) String network analysis of the protein EZHIP in mouse (**B**) and human (**C**). **D**) Bar plot of the log2 difference in the H/L ratio and abundance of the PRC2 complex in oocytes and 2-cell embryos. **E**) Mean phosphosite abundance normalized to the protein abundance of all identified phosphosite on EZHIP in oocytes and 2-cell embryos. The box shows the conserved motif between mouse and human EZHIP. **F, G**) AlphaFold-derived PAE matrices for the hEZHIP complex without (**F**) and with (**G**) phosphorylation within the KLP region. **H, I**) Predicted hEZHIP-hEZH2 interface without (**H**) and with (**I**) phosphorylation of S410 and S412. M406 and R407 are shown in purple and S410/S412 or pS410/pS412 in red. Predicted polar contacts are indicated by yellow dashed lines. hEZH2 is shown as a purple surface and hEZHIP as an orange cartoon. **J**) Introduction of the nomenclature for the peptide sequences of the Biotin-PEG2-conjugated peptides for pull-down experiments. **K**) Western blot of EZH2 on the peptide pull-down experiment with the WT peptide in total HEK cell lysate or crude fractions of cytoplasmatic, organelles, and nuclei fraction. **L**) Box plot of the z-score normalized protein abundances of the PRC2 complex in the peptide pull-downs. Cyt, cytosol; EZH2, Enhancer Of Zeste 2 Polycomb Repressive Complex 2 Subunit; EZHIP, EZH Inhibitory Protein; FDR, false-discover rate; KLP, H3K27M-like Peptide; nuc., nuclear; org., organelle; PAE, Predicted aligned error; PRC2, Polycomb Repressive Complex 2; pSTY, phosphorylation on serine, threonine, and tyrosine; WT, wildtype

Phosphosites exhibiting stable abundance over time included the Cyclin-dependent kinases (CDK) a T14 and CDK12 S1079, both key regulators of the cell cycle and embryonic development (**SI Figure 5D**). Conversely, phosphosites on peptidyl arginine deiminase type VI (PADI6), a core member of the SCMC, displayed divergent behaviour. The protein itself showed slow turnover, whereas the PADI6 S83 phosphosite decreased steadily over the five-day period, suggesting potential regulatory modification of PADI6 function during early development.

### Phosphoproteome and proteome dynamics distinguish oocytes and 2-cell embryos

To investigate how protein synthesis dynamics change during the oocyte-to-embryo transition, protein turnover was compared between oocytes and 2-cell. Despite being developmentally arrested, oocytes exhibited a high degree of protein synthesis, similar to the 2-cell embryos (**Figure 2A**). As observed previously, missed-cleaved tryptic peptides containing lysine residues exhibited a higher fraction of mixed light and heavy SILAC labels compared to arginine-containing peptides indicating higher lysine recycling rates (**Figure 3C** and **Figure 4A**). Overall, oocytes showed a higher degree of amino acid recycling after 24 h compared to 2-cell embryos. Because only a single pSILAC labelling time point was available for this comparison, no kinetic estimation curves were fitted, and heavy-labelled peptide intensities were adjusted by the observed recycling rates.

PCA based on H/L ratios resulted in a clear separation of oocytes and 2-cell embryos (**Figure 4B**). To identify the proteins contributing most strongly to this separation, we examined the corresponding PCA loadings (**Figure 4C**). Whereas the 2-cell embryos were influenced by several proteins, STARD13 and the oocyte-specific linker histone H1FOO (H1F8) were among the strongest drivers of the oocyte grouping. Given the prominent contribution of H1FOO to the separation, we next examined histone synthesis more closely. Direct comparison of H/L ratios between oocytes and 2-cell embryos revealed increased synthesis of most histones in 2-cell embryos, whereas H1FOO showed the opposite pattern, with higher relative synthesis in oocytes (**Figure 4D**). This pattern is consistent with the distinct chromatin requirements of the two stages: mature oocytes are arrested and do not undergo DNA replication, whereas following fertilization, embryos enter rapid cleavage divisions that require extensive synthesis of canonical histones to package newly replicated DNA.[47]

To further dissect differences between protein abundance and synthesis, proteins showing significant changes in either abundance or H/L ratio were plotted against each other (**Figure 4E**). Proteins displaying significant differences were predominantly characterized by high abundance in oocytes but reduced *de novo* synthesis compared to 2-cell embryos. String functional network analysis of these proteins revealed a highly interconnected network enriched for cell-cycle-related regulators, including Wee2, Sall4, Aurka, and Prc1 (**Figure 4F**).[48]

Significant enrichment of GO terms are associated with negative regulation of sister chromatid cohesion, cell cycle checkpoints, chromosome segregation, and transcriptional repressor complexes. These proteins are highly abundant in oocytes but exhibited low synthesis rates, consistent with their degradation during fertilization and early embryogenesis (**Figure 4G**).[49, 50] In contrast, proteins that are both highly abundant and newly synthesized in oocytes were primarily associated with chromatin and gene silencing processes.

Notably, the four proteins NPLOC4, NUP88, LITAF, and CKAP5 displayed low abundance in oocytes but substantially higher synthesis rates compared to 2-cell embryos. Only CKAP5, a microtubule-associated protein, which plays a critical role in spindle assembly and chromosome segregation during mitosis and meiosis, was already identified to be involved in oocyte maturation.[51] Depletion of CKAP5 has been shown to impair oocyte nuclear maturation in both humans and mice, suggesting a role in ovarian aging and early developmental competence.

Given the pronounced changes in the key cell-cycle-regulatory kinases (WEE2 and AURKA) across developmental stages, we next investigated serine, threonine, and tyrosine phosphorylation dynamics. In total, 1,789 phosphosites mapping to 862 proteins were identified. Of these, 249 phosphosites were not annotated in either the Eukaryotic Phosphorylation Site Database (EPSD) or the PhosphoSitePlus database, indicating a substantial number of previously unreported phosphorylation events (**Figure 4H**).[43, 44]

Functional annotation revealed that the identified phosphosites were predominantly associated with processes related to oocyte differentiation and oogenesis, consistent with the enrichment observed for newly identified phosphosites in earlier analyses (**Figure 4I** and **Figure 3P**). Surprisingly, a larger number of phosphosites were quantified in oocytes (∼ 1,600 sites) compared to 2-cell embryos (∼ 1,100 sites) (**Figure 4J**). Although fewer phosphosites were detected on average in 2-cell embryos, PCA showed reduced variability among these samples relative to oocytes (**Figure 4K**). The greater heterogeneity among oocytes may reflect differences in their post-maturation state and associated cellular stress responses, as mature oocytes have limited capacity for prolonged culture and undergo substantial protein degradation during this period.[52–54]

Stage-specific phosphorylation patterns were particularly evident among members of the SCMC. Oocytes showed high phosphosite abundances on several SCMC proteins, including NLRP5 S132, PADI6 S83, ZBED3 S201, and TLE6 T126, which are all proline-directed sites. In contrast, the most abundant phosphosites in 2-cell embryos were associated with proteins involved in protein synthesis and degradation, such as EEF2 T57, EIF4B S497, HUWE1 S3758, and UBE2V2 Y73 (**Figure 4L**).

Reactome pathway enrichment analysis of proteins with significantly altered phosphorylation revealed that 2-cell embryos were enriched for pathways related to cellular communication, including vesicle-mediated transport and membrane trafficking, as well as mitotic anaphase and M-phase progression. Conversely, oocytes showed enrichment for pathways associated with cellular stress responses, including HSF1 activation, and autophagy (**Figure 4M**). Together with the greater variability observed by dimensional reduction, these findings suggest that mature oocytes display a more heterogeneous phosphorylation landscape, potentially reflecting differences in cellular stress and proteome state.

### Phosphorylation in the EZHIP KLP motif modulates its interaction with PRC2

In addition to the identification of mitotic factors, integration of phosphosites and total protein abundance identified a protein involved in chromatin regulation, AU022751, also known as EZH inhibitory protein (EZHIP). Several EZHIP phosphosites were substantially more abundant in oocytes despite relatively modest differences in total EZHIP protein abundance between oocytes and 2-cell embryos (**Figure 5A** and **SI Figure 5F**). String network analyses for both mouse and human, showed an interaction between EZHIP and the protein Enhancer of Zeste Homolog 2 (EZH2), the catalytic subunit of the Polycomb Repressive Complex 2 (PRC2) (**Figure 5B, C**). EZHIP is an inhibitor of PRC2-mediated H3K27 trimethylation (H3K27me3), thereby preventing transcriptional silencing of target genes.[56, 57] PRC2-mediated H3K27me3 regulates the transcriptional repression of key developmental regulators, including homeobox (HBOX) gene clusters and lineage-specifying transcription factors, which are essential for early embryonic patterning and cell fate decisions.[58, 59] Consistent with a role of this regulatory axis in oocytes, the core members of the PRC2 complex (EZH2, SUZ12, EED, RBBP4) were substantially more abundant and showed a higher H/L ratio in oocytes than in 2-cell embryos, suggesting enhanced synthesis for PRC2 in this stage (**Figure 5D**).[60]

Notably, three EZHIP phosphosites were located within or in close proximity to its conserved K27M-like peptide (KLP) motif, a region required for PRC2 inhibition and interaction with EZH2 (**Figure 5E**).[61] Phosphorylation at these sites may therefore modulate EZHIP activity by altering the local interaction interface and reduce or enhance its interaction with EZH2, thereby potentially modulating EZHIP-mediated PRC2 inhibition (**Figure 5F**).

To investigate the structural consequences of phosphorylation, AlphaFold-based predictions were generated for EZHIP and the EZHIP-EZH2 protein complex interactions with and without phosphorylation at S491 and S493 in mouse EZHIP (mEZHIP) and the corresponding sites (S410 and S412) in human EZHIP (hEZHIP). Comparison of the predicted aligned error (PAE) matrices revealed altered prediction confidence for the relative positioning of the EZHIP KLP-containing region upon introduction of the phosphosites in the EZHIP alone or in complex with PRC2 (**Figure 5F, G** and **SI Figure 6A, B, G-J**).[62] While changes in PAE do not directly predict binding affinity, they indicate that phosphorylation affects the confidence of the predicted EZHIP interface. The predicted hEZHIP-hEZH2 interface provided a potential structural explanation for such an effect. In the unmodified model, S410 and S412 in hEZHIP are predicted to build polar interactions, while M406 and R407, residues important for PRC2 inhibition, are positioned within the hEZH2 interaction pocket (**Figure 5H**).[61] Introducing the phosphorylation on hEZHIP at S410 and S412 substantially altered the local electrostatic environment. In particular, the negatively charged phosphate groups introduced an electrostatic repulsion and an additional polar interaction involving R407 resulting in a changed predicted positioning of residues in the KLP motif (**Figure 5I).** A more pronounced structural difference was observed in the corresponding mouse EZHIP-EZH2 models. In the unphosphorylated model, M487 and R488 of mEZHIP were positioned within the predicted mEZH2 interaction pocket, whereas phosphorylation of the serines was associated with displacement of the KLP-containing loop away from the pocket (**SI Figure 6C-F**). For visualization, phosphorylated and unphosphorylated structures were aligned before centering on either model (**Figure 5H, I**). Although these predictions do not establish changes in binding affinity, they support the possibility that phosphorylation alters the local EZHIP-EZH2 interaction interface. These models therefore support the possibility that these phosphorylations perturbs the local EZHIP-EZH2 interaction interface both in mouse and humans.

**Figure 6.**
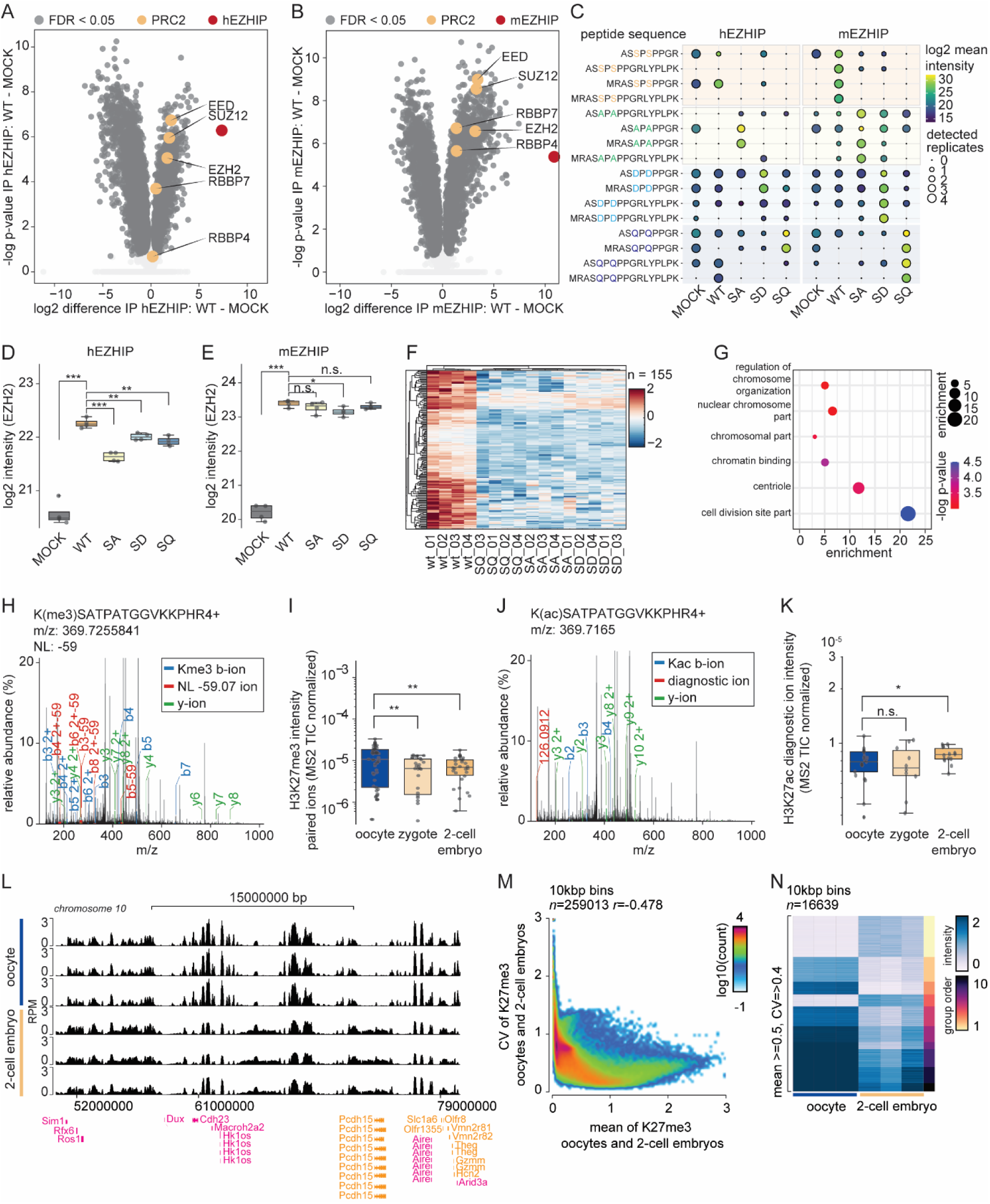
Modification of the EZHIP KLP region modulates PRC2 interaction and coincides with H3K27me3 remodelling during early embryogenesis. A, **B**) Volcano plot of the human EZHIP (**A**) and mouse EZHIP (**B**) IP against the MOCK control. Significantly abundant proteins are highlighted in dark grey, members of the PRC2 complex are highlighted in orange, and the bait in red. Significance was calculated by Student *t*-test, FDR corrected. **C**) Dot plot of identified peptides of the mutated KLP motif in EZHIP. Mutated serine (S) to alanine (A), aspartic acid (D), and glutamine (Q) are highlighted in orange, green, light blue, and dark blue, respectively. Intensity and number of detected replicates are visualized by dot colour and size. **D, E**) Box plot of EZH2 intensity in the human (D) and mouse (E) EZHIP construct IPs. Heatmap of all significantly (*t*-test FDR < 0.05) more abundant proteins in the WT hEZHIP IP compared to the mutants. **G**) Dot plot of selected enriched GO terms from significantly more abundant proteins in the WT hEZHIP IP compared to the mutants. **H**) Representative MS2 spectra of the H3K27me3 peptide precursor 369.7256 m/z. The MS2 TIC for trimethylated (Kme3) fragment b-ions, Kme3 fragment b-ions with the natural loss of 59 m/z, and y-ions are represented in black, blue, red, and green, respectively. **I**) Quantification of the summed paired-intensity of the K27me3 fragment ion normalized to the MS2 TIC, assigned to the H3K27me3 peptide in oocytes, zygotes, and 2-cell embryos. Data is represented in mean ± SEM. Significance is based on Welch’s *t*-test, asterisk represents * p-value < 0.05, ** p-value < 0.01. **J**) Representative MS2 spectra of the H3K27ac peptide precursor 369.7165 m/z. The MS2 TIC for acetylated (Kac) fragment b-ions, the diagnostic secondary acetyl-lysine immonium ion at 126.0913 m/z, and y-ions are represented in black, blue, red, and green, respectively. **K**) Quantification of the Kac diagnostic ion-intensity of the H3K27ac precursor normalized to the MS2 TIC in oocytes, zygotes, and 2-cell embryos. Data is represented in mean ± SEM. Significance is based on Welch’s *t*-test, asterisk represents * p-value < 0.05. **L**) Representative genome browser tracks of H3K27me3 ChIP-seq enrichment in mouse oocytes and 2-cell embryos of Chromosome 10. Y-axis signal is FPKM normalized ChIP-seq read coverage, and genes marked in pink and orange are developmental or lineage specific genes, respectively. **L**) 2D-histogram of mean H3K27me3 signal (X-axis) and the CV of K27me3 (Y-Axis) in oocytes and embryos. Lines represent thresholds used to identify bins with variable signal. **M**) Heatmap of bins with variable H3K27me3 signal in oocytes and embryos. CV, Coefficient of Variation; EED, embryonic ectoderm development; EZH2, Enhancer Of Zeste 2 Polycomb Repressive Complex 2 Subunit; EZHIP, EZH Inhibitory Protein; KLP, H3K27M-like Peptide; PRC2, Polycomb Repressive Complex 2; RBBP4/7, RB Binding Protein 4/7; SUZ12, SUZ12 Polycomb Repressive Complex 2 Subunit; WT, wildtype.

To experimentally test whether modification of the serines in the KLP motif affects PRC2 association, four biotinylated peptides representing the conserved KLP-region of human and mouse EZHIP were synthesized with a PEG2-Biotin tag (**Figure 5L**). The peptides represented the unmodified wild-type sequence (WT), phosphorylation of both serines (pS), serine-to-alanine substitutions (S◊A), and serine-to-glutamine substitutions (S◊Q). As an initial validation, HEK cells were subjected to crude subcellular fractionation and the WT peptide was used for Streptavidin-based affinity enrichment. Western blotting for EZH2 showed the strongest enrichment from the nuclear fraction, whereas little or no EZH2 was detected in the organelle fraction, consistent with the predominantly nuclear localization of PRC2 (**Figure 5K**). The nuclear fraction was subsequently used for Streptavidin-based affinity enrichment with all four EZHIP peptides and unconjugated streptavidin beads as a control, followed by LC-MS/MS analysis. Z-score normalized protein abundances revealed significant enrichment of PRC2 members with the WT peptide, whereas all three modified peptides showed reduced PRC2 complex abundance (**Figure 5M**). The strongest reduction was observed for the phosphorylated and S◊Q peptides. Together, these results demonstrate that alterations of the serines adjacent to the KLP motif reduce association of the EZHIP-derived peptide with PRC2 components and support a phosphorylation-dependent mechanism modifying the EZHIP-PRC2 interaction.

### Serines in the KLP-motif regulate PRC2 association to full-length mouse and human EZHIP

To determine whether the effects observed with EZHIP-derived peptides are maintained in the context of the full-length protein, C-terminally FLAG-tagged human and mouse EZHIP were transiently expressed in HEK cells and immunoprecipitated (IP) using anti-FLAG-antibody beads followed by LC-MS/MS analysis (*n*=4). Both human and mouse WT EZHIP significantly enriched core PRC2 components compared with the MOCK-FLAG control, including EZH2, EED, SUZ12 and RBBP7, as well as RBBP4 for mEZHIP (**Figure 6A, B).** These results confirmed that full-length EZHIP associates with the PRC2 complex under the conditions used for the affinity enrichment.

To investigate the contribution of the two serines adjacent to the KLP motif, the corresponding residues were substituted with alanine (S→A), aspartate (S→D), or glutamine (S→Q). Expression levels of the respective EZHIP variants were confirmed by western blot (**SI Figure 7**) using an anti-FLAG antibody. To further check for the mutated regions in the plasmids, the corresponding peptides were identified and quantified by LC-MS/MS. The expected WT or mutant peptide sequence showed the highest abundance and detection frequency in the corresponding transfection condition (**Figure 6C**). Next, the association of EZHIP with EZH2 was examined in the dependency of the serine substitutions. For hEZHIP, hEZH2 enrichment was significantly reduced for all three mutant variants compared with WT hEZHIP, with the strongest reduction observed for S→A (**Figure 6D**). For mEZHIP, the overall effects were less pronounced, with a significant reduction in hEZH2 enrichment observed for the phospho-mimetic mutation S→D variant (**Figure 6E**). Thus, mutation of the serine-containing region adjacent to the KLP motif alters the interaction of full-length EZHIP with EZH2, although the effect differs between human and mouse EZHIP in transfected human cells.

Because the substitutions affected EZH2 association, the broader EZHIP interactome was investigated for consistent changes. Proteins consistently enriched with WT hEZHIP relative to the mutant variants were identified across the affinity-enrichment experiments. A stringent comparison (*t*-test FDR < 0.05) identified 155 proteins with a higher abundance in the WT EZHIP pull-down (**Figure 6F**). GO analysis of these proteins revealed enrichment for terms associated with chromosome organization, nuclear and chromosomal components, chromatin binding, centrioles, and cell-division sites (**Figure 6G**). Together, these data indicate that alteration of the serine-containing region not only reduces EZH2 association but is accompanied by broader changes in interactions with chromatin- and chromosome-associated proteins.

To determine whether the developmental changes in EZHIP phosphorylation can be associated with changes in PRC2 dependent H3K27me3, we next examined H3K27me3 directly in ArgC digested oocytes, zygotes, and early embryos by LC-MS/MS. The overall trimethylation state was identified based on the identification of the H3K27me3 peptide in the charge states of 2-4. The MS2 scans for the corresponding precursor were used for the quantification of the Kme3-containing fragment ions and the corresponding natural loss (NL) of 59.0735 Da, the trimethylamine-loss, product ions. The signal intensities were paired for the intact and NL fragments and summed across the spectrum. Intensities were normalized to the summed MS2 total ion current (TIC) of the corresponding DIA isolation windows (**Figure 6H**).

Quantification of the paired ion intensity revealed the highest signal in oocytes with a significant reduction to both zygote and 2-cell embryo (p-value = 0.016) (**Figure 6I**). To support this, the diagnostic secondary acetyl-lysine (Kac) immonium ion at 126.0912 m/z from the precursor of H3K27ac (369.7165 m/z) was identified and quantified (**Figure 6J**). The H3K27ac diagnostic ion showed the opposite trend compared to the H3K27me3, increasing from the oocyte to the 2-cell embryo (**Figure 6K**). These results indicate that the oocyte-to-embryo transition is accompanied by a reduction in the global H3K27me3 signal as detected by LC-MS/MS.

To investigate how these developmental differences are reflected across the genome, we next analysed publicly available H3K27me3 ChIP-seq data from mouse oocytes and early embryos.[63, 64] Investigation of normalized H3K27me3 coverage across chromosome 10 revealed extensive differences between oocytes and embryos, including regions of developmental and lineage-associated genes (**Figure 6L**). To systematically identify regions showing pronounced developmental differences, H3K27me3 signal was quantified across 10-kb bins, and regions with variable enrichment were selected based on their mean signal and coefficient of variation (CV) in oocytes and 2-cell embryos (**Figure 6M**). Visualization of the selected regions revealed a broad reduction in H3K27me3 from oocytes to 2-cell embryos (**Figure 6N**), consistent with the difference observed by LC-MS/MS.

Together, these results link developmental regulation of EZHIP to changes in PRC2 association and H3K27me3 during the oocyte-to-embryo transition. Modification of the serine-containing region adjacent to the EZHIP KLP motif reduced its association with EZH2 and altered its broader chromatin-associated interactome, while both LC-MS/MS-based and genome-resolved analyses revealed remodelling of H3K27me3 between oocytes, zygotes and 2-cell embryos. These findings support a model in which phosphorylation-dependent regulation of the EZHIP-PRC2 interaction may contribute to the dynamic control of PRC2 during early embryonic development.

## Discussion

Protein synthesis, rather than mRNA transcript abundance, ultimately executes the developmental program of the early embryo, yet the dynamics of proteome establishment during mammalian embryogenesis have remained essentially unmeasured. Here, we present the first direct, kinetic view of protein synthesis in individual mouse oocytes and early embryos, revealing that the maternally inherited proteome is not a static reservoir but a dynamically replenished and modified system. This fundamentally changes how the maternal-to-zygotic transition should be understood. Early embryonic development is characterized by extensive remodelling of a maternally inherited proteome while developmental control progressively transitions from maternal stored proteins, compounds, and mRNA to embryonic gene expression. Although transcriptomic studies have provided detailed insight into the maternal-to-zygotic transition, changes in transcript abundance do not necessarily reflect changes in protein synthesis, particularly during the earliest developmental stages when stored maternal mRNAs remain an important source for translation.[1, 6] Here, pSILAC enabled us to follow protein synthesis in individual oocytes and early embryos, thereby distinguishing newly synthesized proteins from the abundant pool of maternally deposited proteins. This distinction revealed substantial differences between protein abundance and synthesis that would not be apparent from conventional proteomic measurements. Surprisingly, several components of the SCMC, including NLRP proteins, showed active synthesis during the first 24 h of development despite being considered maternally deposited. Similarly, the cationic amino acid transporters SLC7A1 and SLC7A2, which mediate the uptake of arginine and lysine, were actively synthesized during this period and responded to increased extracellular amino acid concentrations. Together, these observations suggest that proteins required during the earliest developmental stages are not exclusively supplied as a fixed maternal protein pool but can be replenished through translation of maternally deposited mRNA. This is consistent with recent ribosome-profiling studies showing that translation of stored maternal mRNAs continues after fertilization and therefore contribute substantially to the early embryonic proteome.[29]

By extending pSILAC labelling throughout the first five days of development, we were able to follow the progressive transition from a maternally dominated proteome toward an actively synthesized embryonic proteome. This was particularly evident for histones, where the oocyte-specific linker histone H1FOO was one of the major drivers separating oocytes from 2-cell embryos, but most canonical histones showed increased synthesis in 2-cell embryos. This pattern is consistent with the transition from the replication-arrested oocyte to dividing embryos, which require continuous histone synthesis to package newly replicated DNA.[47] Later developmental stages additionally showed increased synthesis of proteins associated with transport and vesicle-mediated processes. Consistent with this observation, newly synthesized proteins were increasingly identified in the culture supernatant, including the lysosomal proteins CTSL, CTSA, and GLB1. Although they all have predicted signal peptides, their secretion path has not been identified yet, and our findings suggest an increased vesicular and lysosomal trafficking as embryos progress through preimplantation development. Interestingly, CTSL has previously been identified in the secretome of high-quality bovine embryos, and supplementation of CTSL to individually cultured bovine embryos improved blastocyst development, hatching, and embryo quality.[45] Together with our detection of newly synthesized CTSL in the mouse embryo supernatant, these observations support further investigation of embryo-derived extracellular proteins as potential mediators or biomarkers of embryo development. Whether similar mechanisms are present in human embryos and could ultimately be informative for non-invasive embryo assessment in assisted reproduction remains to be determined.

In addition to changes in protein synthesis, our data revealed site-specific protein phosphorylation as an important regulatory layer during the oocyte-to-embryo transition. Phosphosite abundances changed independently of the abundance or turnover of the corresponding protein for a set of regulatory proteins. This uncoupling may be particularly important during early development, when rapid cellular reorganization must initially be coordinated using a largely maternally inherited proteome. While protein synthesis and degradation gradually alter proteome composition, phosphorylation provides a mechanism to rapidly modify the state and potentially the function of proteins already present in the oocyte. Consistent with a prominent role for phosphorylation-dependent regulation during this transition, proteins synthesized during the first 24 h were enriched for kinase-associated functions (**Figure 1H**). Moreover, the large number of previously unannotated phosphosites identified in this study indicates that this regulatory layer remains incompletely characterized during early embryogenesis (**Figures 3O** and **4H**).

EZHIP provides a particularly interesting example of such post-translational regulation. Around 30 EZHIP phosphosites showed a higher abundance in oocytes compared to 2-cell embryos and two sites (S491 and S493) were in the conserved KLP motif involved in PRC2 inhibition (**Figure 5A, D**). Structural predictions suggested that phosphorylation could interfere with the local interaction site between EZHIP and EZH2, while peptide pull-down experiments showed reduced PRC2 association following modification of the serines in the KLP-motif. Importantly, this effect was reproduced in the context of full-length EZHIP. Substitution of the serines reduced EZH2 abundance in the pull-down, particularly for human EZHIP, and showed broader changes in the EZHIP interactome. Proteins pulled down with WT EZHIP were enriched for chromatin- and chromosome-associated functions, further connecting this region of EZHIP to chromatin regulation.

Regulation through PTMs may be particularly relevant during the oocyte-to-embryo transition, when the H3K27me3 landscape is extensively reorganized. Consistent with this, LC-MS/MS-based analysis of the H3K27me3 peptide revealed decreasing H3K27me3 signal from oocytes to zygotes and 2-cell embryos, while analysis of published H3K27me3 ChIP-seq data demonstrated extensive remodelling genome wide of H3K27me3 between oocytes and 2-cell embryos. Recent studies have independently established an important role for maternal EZHIP during this developmental period, showing that EZHIP translated from maternal mRNA restricts PRC2 enzymatic activity during the first embryonic divisions and contributes to the regulation and inheritance of H3K27me3-dependent chromatin states.[56, 65, 66] Together, our findings suggest that regulation of EZHIP during early development may extend beyond its abundance and include PTM regulation of its PRC2-interacting region. Phosphorylation or dephosphorylation could thereby provide a rapid cellular mechanism to modulate EZHIP-PRC2 association during developmental stages characterized by epigenetic reprogramming. However, whether the identified phosphorylation events directly contribute to the observed developmental changes in H3K27me3 remains to be established.

Beyond early embryonic development, phosphorylation-dependent regulation of EZHIP may also be relevant in the oncology. EZHIP is normally associated with germline and developmental contexts but is aberrantly highly expressed in posterior fossa group A ependymomas and a subset of H3K27-altered diffuse midline gliomas, where inhibition of PRC2 contributes to the characteristic reduction in H3K27me3.[67] Consequently, the identification of phosphorylation as a potential regulator of the EZHIP-PRC2 interaction raises the possibility that the phosphorylation state of EZHIP could influence its inhibitory activity also in EZHIP-driven tumours. Determining the kinases and phosphatases controlling these sites, and whether modulation of EZHIP phosphorylation is sufficient to alter PRC2 activity and H3K27me3 in cellular models, will therefore be important directions for future studies.

Finally, the regulation of EZHIP and PRC2 may also be relevant for the rapidly developing field of stem-cell-based embryo models. These systems increasingly recapitulate key morphological, transcriptional, and cellular features of early embryogenesis and provide experimentally accessible models for developmental processes that are difficult to investigate in embryos. However, achieving molecular and epigenetic vitality to the embryo remains an important challenge. Our findings, together with the emerging role of maternal EZHIP in restricting PRC2 activity and controlling H3K27me3 reprogramming during the first embryonic divisions, highlight the importance of considering post-translational and chromatin-regulatory mechanisms when establishing and evaluating *in vitro* models. Consequently, the developmental dynamics of EZHIP phosphorylation, PRC2 activity, and H3K27me3 could provide additional molecular markers against which stem cell-based embryo models can be compared with the *in vivo* model. Integrating LC-MS/MS-based (phospho-)proteome analysis next to transcriptomic and epigenetic profiles could help determine how accurately these models replicate the regulatory transitions that occur during early embryogenesis.

Several limitations should be considered in our study. Heavy SILAC amino-acid incorporation was relatively low in oocytes and during the first 48 h of the embryo development because of the substantial contribution of recycled and maternally derived light amino acids. Correction for amino-acid recycling is necessary for calculating protein synthesis and turnover. Furthermore, the study was performed in mouse oocytes and embryos cultured *in vitro.* While the embryos developed morphologically normal, differences cannot be excluded. Furthermore, to count for biological differences, single oocytes and embryos were analysed, limiting the material used for the phosphoproteomic analysis. While achieving here a considerable deep phosphoproteome for single-cells and embryos, phosphosite coverage remains incomplete. Most importantly, although structural modelling, peptide pull-downs, and full-length EZHIP affinity enrichment consistently support a role for the serines in the KLP-motif in regulating PRC2 inhibition, the functional consequences of their phosphorylation have not been directly tested in embryos. The observed developmental changes in EZHIP phosphorylation and H3K27me3 therefore remain correlative. Targeted mutation of these phosphosites in oocytes or embryos will be required to establish whether EZHIP phosphorylation directly controls PRC2 activity and H3K27me3 reprogramming *in vivo*.

In conclusion, our findings extend previous transcriptomic and ribosome-profiling studies by providing a proteome-resolved view of protein synthesis and turnover during early embryogenesis. Building on recent advances in single-cell proteomics [28], we demonstrate that pSILAC can be applied at the single-embryo level to distinguish newly synthesized proteins from the maternally inherited proteome. We demonstrate that the maternal proteome is not a static proteome but is continuously renewed, recycled, and post-translationally regulated during the oocyte-to-embryo transition, providing multiple layers of regulation through which early embryos reorganize their proteome and chromatin landscape before and during ZGA.

## Materials and Methods

### Animals

Animal work was conducted according to license no. 2021-15-0201-00851, approved by the Danish National Animal Experiments Inspectorate, and performed according to national and local guidelines. Mice were kept in designated rooms in individually ventilated cages at a temperature of 22 °C (± 2 °C), with a humidity of 55% (± 10%), air in the room was changed eight to ten times per h and dark/light cycle is 12 h/12 h, light from 6 am to 6 pm. Mouse husbandry is performed according to Danish regulations for animal experiments.

### Isolation of mouse oocytes and zygotes

Ovulation of prepubescent (4-week old) C57BL/6NRj females (mus musculus) (*n*=41) was induced by an intraperitoneal (IP) injection of PMSG (HOR-272, Prospec), 5 IU/female, followed by an IP injection of hCG (Chorulon Vet, Pharmacy), 5 IU/female, 47 h later. After the second injection, the female mice were mated with C57BL/6NRj stud breeding males. The next morning, the females were euthanized and the oviducts were dissected to harvest the cumuli containing zygotes and unfertilized oocytes. Cumuli were disaggregated by incubating in Hyaluronidase (H4272, Sigma-Aldrich) for 10 min. Degraded oocytes were excluded from the experiments. Zygotes were sorted out assessed by the presence of the second polar body and cultured in KSOM medium supplemented with essential amino acids (50*X*) as detailed in **Table 1** and non-essential amino acids except L-lysine and L-arginine.

**Table 1.** Essential amino acids composition for 50X medium.

| Amino Acid | Molecular Weight | Concentration (mg/L) | mM | Medium abbreviation |
| --- | --- | --- | --- | --- |
| [<sup>12</sup>C<sub>6</sub>, <sup>14</sup>N<sub>4</sub>]-L-arginine | 211 | 6320 | 30 | 1X light (1X L) |
| [<sup>13</sup>C<sub>6</sub>, <sup>15</sup>N<sub>4</sub>]-L-arginine | 221 | 6607 | 30 | 1X light (1X H) |
| [<sup>12</sup>C<sub>6</sub>, <sup>14</sup>N<sub>4</sub>]-L-arginine | 211 | 31,600 | 150 | 5X light (1X L) |
| [<sup>13</sup>C<sub>6</sub>, <sup>15</sup>N<sub>4</sub>]-L-arginine | 221 | 33,035 | 150 | 5X light (1X H) |
| L-Cystine | 240 | 1200 | 5 |  |
| L-Histidine hydrochloride-H<sub>2</sub>O | 210 | 2100 | 10 |  |
| L-Isoleucine | 131 | 2620 | 20 |  |
| L-Leucine | 131 | 2620 | 20 |  |
| [<sup>12</sup>C<sub>6</sub>, <sup>14</sup>N<sub>2</sub>]-L-lysine | 183 | 3625 | 3.62<br>5 | 1X light (1X L) |
| [<sup>13</sup>C<sub>6</sub>, <sup>15</sup>N<sub>2</sub>]-L-lysine | 191 | 3775 | 3.62<br>5 | 1X light (1X H) |
| [<sup>12</sup>C<sub>6</sub>, <sup>14</sup>N<sub>2</sub>]-L-lysine | 183 | 18125 | 18.1<br>25 | 5X light (1X L) |
| [<sup>13</sup>C<sub>6</sub>, <sup>15</sup>N<sub>2</sub>]-L-lysine | 191 | 18,875 | 18.1<br>25 | 5X light (1X H) |
| L-Methionine | 149 | 755 | 5.06 |  |
| L-Phenylalanine | 165 | 1650 | 10 |  |
| L-Threonine | 119 | 2380 | 20 |  |
| L-Tryptophan | 204 | 510 | 2.5 |  |
| L-Tyrosine | 181 | 1800 | 1.8 |  |
| L-Valine | 117 | 2340 | 20 |  |

For light medium, [¹²C₆, ¹⁴N₂]-L-lysine and [¹²C₆, ¹⁴N₄]-L-arginine were added, while the heavy medium contained [¹³C₆, ¹⁵N₂]-L-lysine and [¹³C₆, ¹⁵N₄]-L-arginine On the day of harvesting a group of both zygotes and oocytes were processed for analysis, while other zygotes were incubated in KSOM at 37 °C and 5% CO2 for up to 5 days. At the different developmental stages, oocytes and embryos were washed in PBS (20012-027, Life Technologies). and pippeted with a 50 μm diameter glass capillary in PBS afterwards.

### EmbryoScope

Videos of the development from the embryos were uploaded to 10.5281/zenodo.22876670. The 16 microwells in the EmbryoSlides (Vitrolife, Denver, CO), each with a diameter of approximately 250 μm, were filled according to the manufacturer’s instructions with the specific medium designated for oocytes or zygotes as described above. The microwells and wells were overlaid with 1.6 mL of mineral oil (Sigma-Aldrich, St. Louis, MO) and equilibrated in an incubator at 37 °C in a humidified atmosphere of 5% CO2 in air.

Oocytes or zygotes were loaded into the wells of the EmbryoSlide containing pre-equilibrated medium. EmbryoSlides were then loaded into the EmbryoScope + ™. Oocytes or zygotes were cultured for 24-144 h at 37 °C in a humidified atmosphere of 5% CO2 in air. Images were taken every 10 min at 11 focal planes with low-intensity red LED illumination with < 0.5 s of light exposure per image. These conditions are identical to those used for humans in ART, and therefore, are considered to have minimal impact (if any) on gametes and preimplantation embryos.

### Cell culture

HeLa human cervix carcinoma cells (ATCC CCL-2) and human embryonic kidney (HEK) cells were cultured in DMEM (Gibco, Invitrogen), supplemented with 10% fetal bovine serum, 100 U/ml penicillin (Invitrogen), 100 μg/ml streptomycin (Invitrogen), at 37 °C, in a humidified incubator with 5% CO2. At ∼80% confluence, cells were detached using trypsin and washed twice with Phosphate Buffered Saline (PBS) from Gibco (Life Technologies).

### Single-cell and low-input digestion

Isolated oocytes, embryos, or cells were digested following the One-Tip [21] or the single-cell in 96-well plate [25] protocol. In brief, for the OneTip protocol, Evotips were pre-conditioned with 20 µl buffer B composing of 100% Acetonitrile (ACN), 0.1% formic acid (FA) and centrifuged at 800 xg, 55 s. After 10 s activation with isopropanol, tips were washed with 20 µl buffer A (0.1% FA) and centrifuged. 2.5 µl of lysis and digestion buffer 1 (0.2% DDM, 100 mM TEAB, 20 ng/µl Trypsin, 10 ng/µl Lys-C) or 2 (0.05% DDM, 30 mM TEAB, 5 ng/µl Trypsin, 2.5 ng/µl Lys-C, 0.25 mM DTT) was added and spun down at 50 xg for 30 s. Samples (n>4) were loaded on the tips and incubated at 37 C for 15 min or 3 h. Digestion was stopped by washing the tips two times with buffer A. The in plate digestion was performed for ArgC digestions of single-cell or low-input material. In the 96-well plate, 3 µl of digestion buffer 3 (0.05% DDM, 30 mM TEAB, 10 ng/µl ArgC, 0.25 mM DTT) was placed and oocyte, zygotes, 2-cell embryos, or ∼2000 cells were added into the plate in a maximum volume of 7.5 µl. Plate was centrifuged for 20 sec, 400 xg to collect cells at the bottom of the well. Digestion was performed for 3h at 37 °C. Digestion was stopped by acidifying with 2 µl 5% FA in 2% ACN.

### Western Blot

Lysed cells were incubated with NuPAGE™ LDS Sample Buffer (4X) (ThermoFisher Scientific) with 5 mM DTT for 10 min at 95 °C. NuPAGE Bis-Tris gels (Invitrogen) were run according to the manufacturer’s instructions at 120 V in MOPS buffer. Gels were washed in Transfer buffer (25 mM Tris, 192 mM glycine, 10% ethanol). PVDF membranes were activated with methanol and proteins were transferred in transfer buffer at 20 V over 2 h. Membranes were blocked with 5% milk in PBS-T and incubated with 1:2000 EZH2 (Millipore) or 1:2000 anti-FLAG-HRP (Merck) over night at 4 °C. Membranes were washed 3x with PBS-T for 10 min. Anti-rabbit secondary antibody was incubated for 1 h at RT. After washing, membranes were developed with Pierce ECL Western Blotting Substrate (ThermoFisher Scientific).

### Peptide Pull-down preparation

HEK cells were cultivated on 15 cm plates and washed 2x with 4 °C cold PBS. Cells were scraped of in 5 mL 4 °C cold PBS and centrifuged at 300 xg for 3 min at 4 °C. Cells were splitted 3:1 and subjected to crude fractionation or total cell lysis. Crude fractionation was achieved by resuspending the cell pellet in 950 µl hypotonic homogenization buffer (25 mM Tris-HCl pH 7.5, 50 mM sucrose, 0.2 mM EGTA, 0.5 mM MgCl2) and mechanically homogenized by passing the cell suspension 4 times through a 23G blunt 1” needle (SAI Infusion Technology #B23-100) attached to a 1 mL syringe (Air-Tite NormJ-ect-F #NJ-9166017-02). Directly, 89 µl of concentrated sucrose buffer (2.5 M sucrose, 0.2 mM EGTA, 0.5 mM MgCl2) were added to restore tonicity. Samples were centrifuged at 1,000 ×g for 10 min (4°C) and cell pellet, which should contain mainly nuclei, and total cells were lysed with solubilization buffer (1% Triton x-100, 150 mM NaCl, 10 mM KCl, 50 mM Tris pH8.5, 5 mM MgCl2, Benzonase, with protease inhibitor and phosphatase inhibitors). Supernatant was further centrifuged at 20,000 xg for 30 min (4°C) to collect the organelle fraction in the pellet and the cytoplasmic fraction in the supernatant. Organelle fraction was lysed with solubilization buffer and 100 µl of 10X solubilization buffer was added to the cytoplasmic fraction. Total input, nuclei, organelles, and cytoplasm were rotated for 30 min in a head-over-head rotation wheel at 4 °C. Protein lysates were cleared by centrifugation for 15 min at 15,000 xg at 4 °C. Aliquots were taken for proteome analysis.

Pierce™ Streptavidin Magnetic Beads (Thermo Scientific™) were prepared by washing 2x with PBS-T and resuspending the beads in 7 mL of 4% Formaldehyde, followed by 7 mL of 0.2 M NaCnBrH and incubation for 2 h at RT. Supernatant was removed and washed with 10 mL of 0.1 M Tris-HCl (pH 7.5), and 2x with PBS-T. Beads were resuspended in 5 mL of PBS-T after washing and kept at 4 °C until further use.

Modified Streptavidin Magnetic Beads were washed 2x with solubilization buffer and incubated with an excess of peptide (1:2 beads:peptides ratio) for 2 h at RT. Conjugated beads were washed 2x with solubilization buffer and added to the protein lysates from the total, nuclei, organelle, and cytoplasmic input.

### Plasmid pull-down preparation

HEK cells were transfected by a calcium-phosphate transfection. In brief, 4 h before the transfection, 0.3 x 10^6 HEK cells were plated out into a 6-well plate. DNA (2 µg) was diluted in 67.5 µl sterile water and 15 µl 2.2 M CaCl2. DNA solution was added dropwise into 75 µl 2X HBS (50 mM HEPES, 280 mM NaCl, and 1.42 mM Na₂HPO₄ (pH 7.05)) and mixed well, before 20 min incubation at RT. The transfection solution was added dropwise to the cells and incubated for 48 h.

Cells washed 2x with 4 °C cold PBS, lysed by adding and scraping off in 150 µl modified RIPA (mRIPA) buffer (1% Triton x-100, 50 mM Tris-HCl (pH 7.5), 150 mM NaCl, with protease and phosphatase inhibitors), and were rotated for 30 min in a head-over-head rotation wheel at 4 °C. Protein lysates were cleared by centrifugation for 15 min at 15,000 xg at 4 °C. Aliquots were taken for proteome analysis.

Pierce™ Anti-DYKDDDDK Magnetic Agarose was washed 2x with mRIPA buffer and added to the protein lysates.

### Automated plasmid and peptide pull-down

Peptide and plasmid pull-down were performed using the KingFisherTM Flex robot (Thermo Fisher Scientific). Protein lysates with beads were adjusted to the total volume of 500 µl with mRIPA or solubilization buffer. The robot was set up as the followed: the 96-well comb is stored in plate #1, sample plate #2. Washes were performed in the plates #3-#7 with 2x solubilization buffer, 2x 150 mM NaCl, and 1x MilliQ water. Beads were released into 100 µl 50 mM TEAB, followed by reduction and alkylation with the final concentrations of 5 mM TCEP and 15 mM CAA.

### Automated bulk proteome digestion

Bulk proteome was reduced and alkylated by adding to a final concentration of 5 mM TCEP and 10 mM CAA and incubating for 20 min at RT. Afterwards, samples were digested overnight using the PAC protocol [68] implemented for the KingFisher^TM^ Flex robot (Thermo Fisher Scientific) in 96-well format [69, 70] as described previously. Protein lysates were adjusted to a total of 300 µl with lysis buffer. The 96-well comb is stored in plate #1, the sample in plate #2 in a final concentration of 70% acetonitrile and with 5 µl of magnetic Amine beads (ReSyn Biosciences) in a protein/bead ratio of 1:2. Washing solutions are in plates #3–5 (95% ACN) and plates #6–7 (70% Ethanol). Plate #8 contains 100 μl digestion solution of 50 mM TEAB, 0.25 µg of LysC (KPL) and 0.5 µg trypsin (Sigma Aldrich). Protein aggregation was carried out in two steps of 1 min mixing at medium mixing speed, followed by a 10 min pause each. The sequential washes were performed in 2.5 min and slow speed, without releasing the beads from the magnet. Beads were released into the digestion buffer and incubated at 27 °C overnight. Samples were acidified after digestion to final concentration of 1% FA. 5 µl of each sample were loaded directly into Evotips (Evosep) for full proteome analysis.

### LC-MS/MS setup for SILAC samples

Peptide separation was performed on an Evosep ONE or Evosep ENO system (both Evosep, Denmark) equipped with an Aurora® Elite™ 15×75 C18 UHPLC column (IonOpticks, Australia) with the preprogrammed 40 (31 min) samples per day (SPD) whisper method.

Mobile phases consisted of 0.1% FA as solvent A and 0.1% FA in ACN as solvent B. The HPLC system was coupled to an Orbitrap Astral mass spectrometer or Orbitrap Astral Zoom mass spectrometer using Nanospray Flex ion source (Thermo Fisher Scientific).

For DDA experiments, the Orbitrap Astral mass spectrometer was operated with a fixed cycle time of 0.5 s and with a full scan range of 380-980 m/z at a resolution of 180,000. The automatic gain control (AGC) was set to 500%. Precursor ion selection width was kept at 2-Th and peptide fragmentation was achieved by HCD (Normalized Collision Energy 30%). Fragment ion scans were recorded at a resolution of 80,000 and maximum fill time of 2.5 ms. Dynamic exclusion was enabled and set to 10 s.

For DIA experiments, the Orbitrap Astral (Zoom) mass spectrometer was operated at a full-MS resolution of 240,000 with a full scan range of 380-980 m/z when stated. The full-MS AGC was set at 500%. For the Astral mass spectrometer, fragment ion scans were recorded at a resolution of 80,000, isolation window of 4 Th, and maxIT of 6 ms. The Astral Zoom mass spectrometer recorded MS2 scans with an isolation window of 4 Th and maxIT of 4.4 ms. The isolated ions were fragmented using HCD with 25% Normalized Collision Energy (Guzman et al. 2024).

### LC-MS/MS setup for IPs and ArgC digested samples

Peptide separation was performed on an Evosep ENO system (both Evosep, Denmark) equipped with an Pepsep 8 cm column for peptide pull-downs and an Aurora® Elite™ 5×75 C18 UHPLC column (IonOpticks, Australia) for plasmid pull-downs with the preprogrammed 100 SPD (13 min) method and 80 (16 min) SPD whisper method. For ArgC digested samples, the Vanquish NEO was used for linear peptide separation operating on 80% ACN, 0.1% FA for the organic phase and 0.1% FA for buffer A. The HPLC systems were coupled to an Orbitrap Astral Zoom mass spectrometer using Nanospray Flex ion source (Thermo Fisher Scientific).

The mass spectrometer was operated at MS1 resolution of 240,000 at a scan range of 380-980 and 350-1050 m/z. The fragment ion scans were recorded from a scan range of 150-2000 m/z or 125-1800 m/z with the isolation windows of 2 Th for a maxIT of 1.5 or 2 ms.

### Data analysis

Code is available via Zenodo with the DOI 10.5281/zenodo.22878819. All MS runs were searched with DIA-NN v2.2.0. For human and mice in silico generated spectral library were generated from a homo sapiens and murine protein sequence fasta downloaded from Uniprot downloaded in April, 2024. For standard SILAC search, SILAC,0.0,KR was used as fixed modification with the channels of SILAC,L,KR,0:0 and SILAC,H,KR,8.014199:10.008269. Additional parameters were specified to Mass accuracy of 15 ppm, and MS1 accuracy of 5 ppm. Normalization was performed with “channel-run-norm“ and files were searched with match between runs. To identify phospho sites and to calculate the recycling rate, two in silico generated spectral library were generated with heavy lysine (SILAC_K,8.014199,K) and heavy arginine (SILAC_R,10.008269,R) as variable modifications, or “Phospho” (Steger et al. 2021) as variable modification with two missed cleaves and maximum of five variable modifications for mice.

### Protein Quantification and SILAC incorporation

DIA-NN reports were imported into R v4.3.3 and processed using the packages dplyr, tidyr, stringr, tibble, matrixStats, reshape2, arrow, and ggplot2. Precursors were filtered at Lib.PG.Q.Value < 0.01 and Translated.Q.Value < 0.01.

For each run, precursor or protein intensities were separated according to the light (L) and heavy (H) SILAC channels and aggregated to the peptide and protein levels. Heavy-label incorporation was calculated as: Incorporation = H/(H+L). The H/L ratio was calculated as: Ration = H/L. Proteins detected in only one channel were used only for the abundance of the protein, which was calculated as: Abundance = H + L.

For analyses based on relative intensity-based absolute quantification (riBAQ), iBAQ values were divided by the summed total iBAQ value for the respective sample.

### Recycling and turnover

DIA-NN parquet reports (Recycle with SILAC as variable, and Bulk with SILAC as Channel) were imported into R v4.3.3 and processed using the packages dplyr, tidyr, stringr, tibble, matrixStats, reshape2, arrow, and ggplot2. Precursors were filtered at Lib.PG.Q.Value < 0.01 and Translated.Q.Value < 0.01.

To estimate recycling of unlabeled lysine and arginine, a separate variable-modification SILAC search (“Recycle” search) was used. Recycling was estimated independently for lysine and arginine. Peptides containing exactly two residues of the respective amino acid were retained, allowing three labeling states to be distinguished based on the number of SILAC modifications in the precursor identifier: pepLL, containing two light amino acids; pepLH, containing one light and one heavy amino acid; and pepHH, containing two heavy amino acids. MS1 areas were averaged for each peptide precursor and run. Peptides detected in both the LH and HH states were retained for estimation of heavy-amino-acid availability, and the corresponding intensity matrices were aligned across peptides and runs.

Assuming independent incorporation of heavy amino acids at the two labeling positions, the expected fractions of the three labeling states are

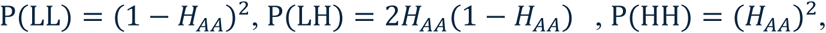

where *H_AA_* represents the fraction of the intracellular lysine or arginine pool available in heavy form. Heavy-amino-acid availability was estimated from the relative intensities of LH and HH peptide species as 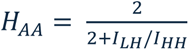.

The median peptide-level estimate was calculated for each run. Accordingly, values approaching 1 represent predominantly heavy amino-acid availability, whereas lower values indicate an increasing contribution of recycled light amino acids.

The time dependence of heavy-amino-acid availability was modelled separately for lysine and arginine using a biexponential function: *H_AA_*(*t*) = 1 − [*ce*^−*at*^ + (1 − *ce*^−*bt*^)].

where *a* and *b* describe the fast and slow components of heavy-amino-acid incorporation and *c* their relative contribution. The curves were anchored at *H_AA_*(0) = 0, and parameters were estimated by nonlinear least-squares fitting. If the three-parameter biexponential model did not converge, a biexponential model with (*c* =0.5) was used, followed by a single-exponential model as an additional fallback.

For correction of the main SILAC dataset (“Main” search), peptides were separated according to their C-terminal lysine or arginine and processed independently using the corresponding amino-acid-specific recycling curve. In contrast to the recycling estimation described above, peptides in the Main search were not required to contain two lysine or arginine residues. Light and heavy MS1 areas were aggregated for each peptide precursor and run.

For peptide/run combinations with sufficient light and heavy signal, an apparent peptide-specific turnover rate α was numerically estimated from the measured light intensity, total peptide intensity, labelling duration, and the experimentally determined amino-acid recycling kinetics. The corresponding fraction of newly synthesized peptide was calculated as

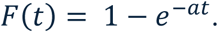

Where direct estimation of α was not possible because one labelling state was not detected, *F*(*t*) was inferred from the available light or heavy peptide signal together with the predicted heavy-amino-acid availability at the corresponding labeling time. Estimated fractions were constrained between 0 and 1.

The resulting peptide-specific newly synthesized fractions and the lysine- or arginine-specific *H_AA_*(*t*) values were used to calculate recycling-adjusted heavy peptide intensities. In the final analysis, measured light intensities were retained, whereas heavy intensities were adjusted according to the estimated newly synthesized fraction and the predicted heavy-amino-acid availability at the corresponding labeling time. Thus, recycling correction was performed separately for lysine and arginine and accounted for the time-dependent availability of heavy amino acids rather than applying a single global correction factor.

Following recycling correction, lysine- and arginine-terminated peptides were combined for protein-level quantification. Peptides were ranked within each protein based on their mean total intensity after run-wise median normalization, and up to the three most intense peptides per protein were retained. Light and heavy protein intensities were calculated as the median intensity across the selected peptides for each run. Total protein intensity, heavy-label incorporation, and the H/L ratio were calculated as described above.

For the developmental time-course analysis, recycling-corrected incorporation profiles were used for subsequent estimation of apparent protein turnover and protein half-lives. For comparisons for which only a single labelling time point was available, including the comparison between oocytes and 2-cell embryos, no kinetic turnover curves were fitted; instead, SILAC incorporation was corrected using the recycling estimate corresponding to the respective labelling period.

### Targeted Trimethylation and Acetylation analysis

Raw files were converted to mzML files via MSconvert (ProteoWizard 3.0.25035) and used as input for a python (v3. 9) based analysis using anaconda prompt. Python scripts are available on Zenodo 10.5281/zenodo.22878819. The ArgC-digested peptide K(me3/ac)SAPATGGVKKPHR were monitored at their theoretical precursor charge states 369.7255841, 492.6316867, 738.4438917. Because of the low absolute mass difference between the PTMs of trimethylation ((-C₃H₆) +42.0470 Da) and acetylation ((-C₂H₂O) +42.0106 Da), modifications were analysed by either the neutral loss of −59 Da of trimethylation or by their respective diagnostic ion at m/z 126.0913. Even though trimethylation retains and acetylation neutralizes the positive charge of lysine, the charge states were not accounted for potential biases. MS2 spectra from DIA isolation windows containing the respective precursors were searched with a mass tolerance of 10 ppm for theoretical sequence fragments and the characteristic 59.0735 Da NL of trimethylamine from Kme3-containing fragment ions.

For each spectrum, Kme3-containing fragment ions and their corresponding neutral-loss products were quantified. A fragment pair was defined by co-detection of a theoretical Kme3-containing fragment and its corresponding trimethylamine-loss ion. Spectra containing at least two such fragment pairs together with at least one additional sequence-supporting fragment ion were considered to provide supporting evidence for the modified peptide. Fragment-pair intensities were calculated using the lower intensity of the intact and neutral-loss fragment for each pair and summed across spectra. Signal intensities were additionally normalized to the summed MS2 total ion current (TIC) of the corresponding DIA isolation windows. Retention time was recorded for inspection but was not used for filtering or scoring.

Acetylation of H3K27 was monitored by the diagnostic ion m/z 126.0913 in the above-mentioned theoretical precursor charge states.

### ChiP-seq data analysis

Previously published H3K27me3 ChIP-seq datasets from fully grown oocytes, 2-cell, and 8-cell embryos [64] were re-analysed using a processing pipeline described previously by Halliwell et al.[63] Briefly, sequencing quality was assessed using FastQC [71] and FastQ Screen [72], reads were trimmed with Trim Galore!, aligned to the mm10 reference genome using Bowtie2 [73], and processed using SAMtools [74]. Quality-control summaries were compiled with MultiQC [75]. Downstream quantification and visualisation were performed in EaSeq [76]. H3K27me3 enrichment was quantified in 10-kb genomic bins, and regions showing developmental remodelling were identified based on signal intensity and coefficient of variation between oocytes, 2-cell, and 8-cell embryos. Heatmaps, two-dimensional histograms, and genome browser tracks were generated in EaSeq.

### Statistical Analysis

Statistical analyses were performed in R v4.3.3 or Perseus v1.6.15.0. Unless otherwise indicated, data are presented as individual biological replicates together with the corresponding group summary statistics. For comparisons between experimental groups, statistical significance was assessed using two-sided Student’s t-test or Welch’s t-test, as indicated in the corresponding figure legends. For proteome- and phosphoproteome-wide comparisons, multiple-hypothesis testing was controlled using the Benjamini–Hochberg procedure or permutation-based FDR, as specified for the respective analysis. Proteins or phosphosites were considered significantly altered at FDR < 0.05. s0 = 0.1.

Principal component analysis (PCA) was performed on log2-transformed protein intensities and H/L ratios. Only proteins quantified with a 70% data completeness were included. PCA loadings were used to identify proteins contributing most strongly to the separation of experimental groups.

For temporal analyses, protein synthesis profiles were clusters using k-means, and functional enrichment analyses were subsequently performed for individual clusters. GO and pathway enrichment analyses were evaluated using gprofiler or Fisher’s Exact test. Enrichment P values were corrected for multiple testing using the Benjamini-Hochberg procedure, with an adjusted P value < 0.05 considered significant.

**SI Figure 1.**
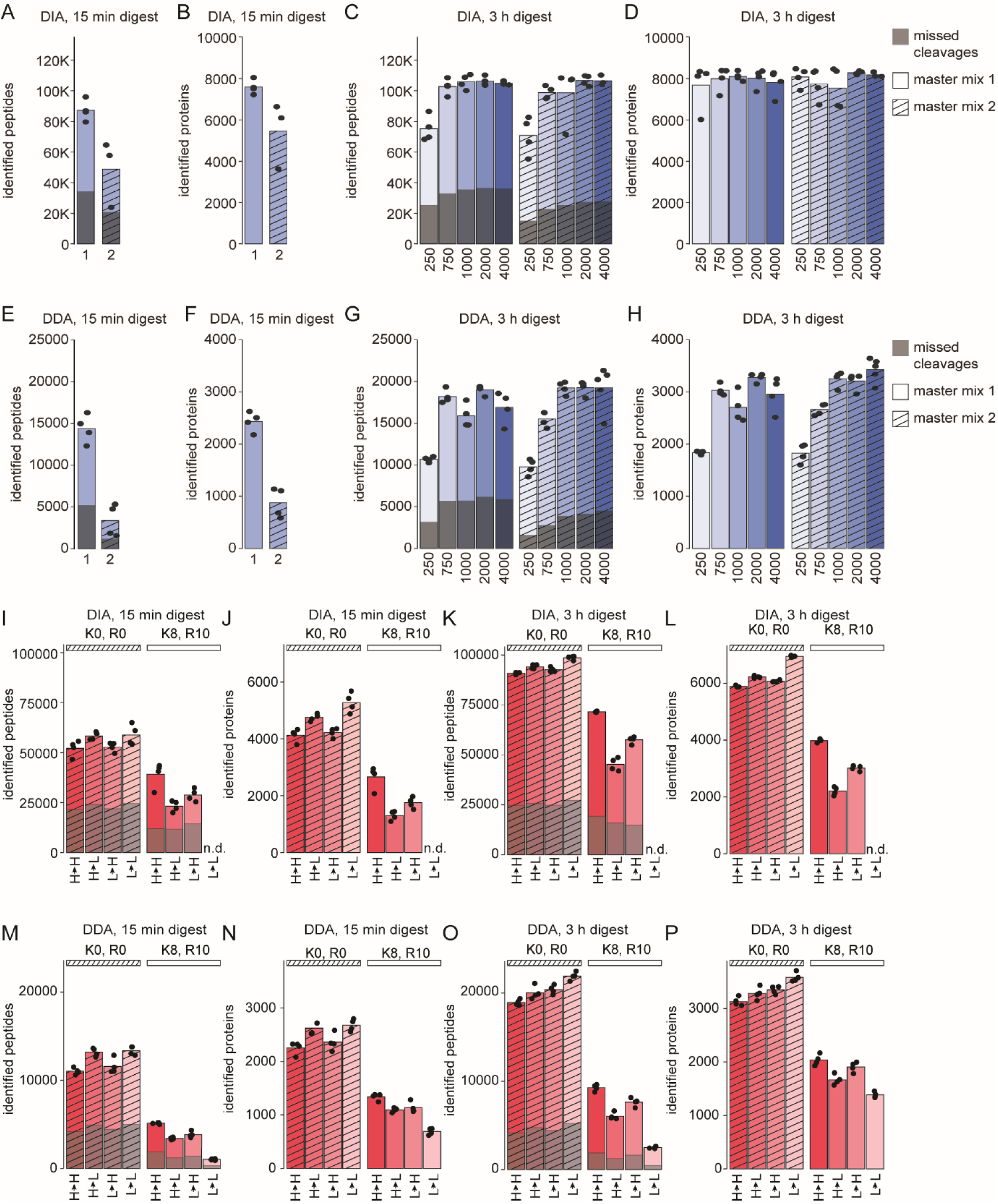
**Optimal digestion conditions and acquisition strategies. A-H**) Comparison of two digestion and lysis master mixes based on peptide and protein identifications obtained using 15 min (**A, B, E, F**) or 3 h (**C, D, G, H**) digestion times. Master mix 2 is shown striped. **I-P**) Comparison of the effects of short and long digestion times and DIA and DDA analyses in pSILAC-treated HeLa cells. Identified peptides and proteins from both light and heavy isotope channels are presented as bar plots for 15 min (**I, J, M, N**) and 3 h (**K, L, O, P**) digestion times for DIA (**I-L**) and DDA (**M-P**). The light channels are shown striped.

**SI Figure 2.**
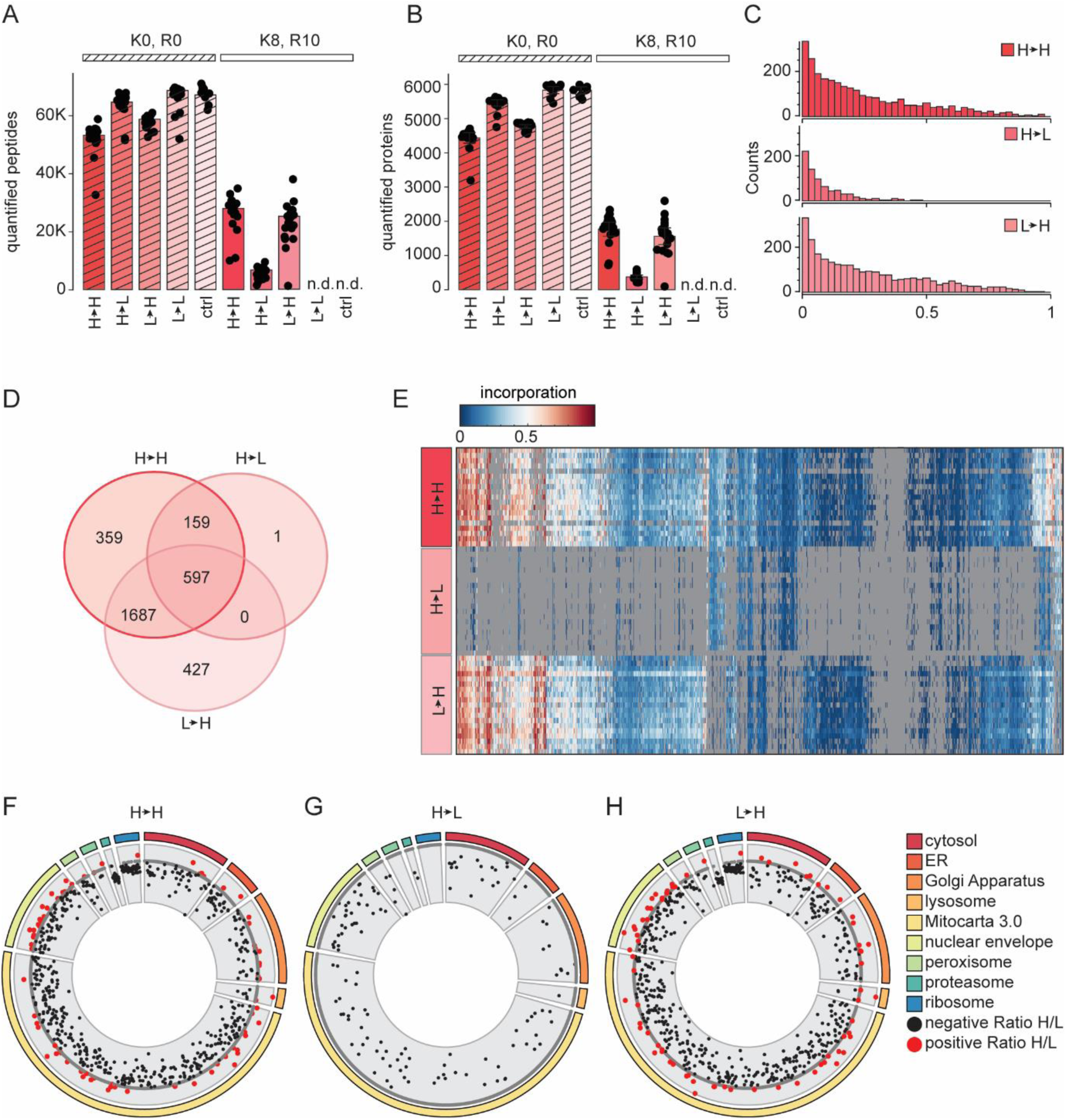
Metabolic labelling of zygotes over 48 h. A, **B**) Identified peptides (**A**) and proteins (**B**) in the light and heavy SILAC channels using DIA-NN for data analysis. The light channels are shown striped. **C**) Histogram of incorporation levels per normalized counts across the three conditions H◊H, H◊L, and L→H. **F, G**) Venn diagram (**F**) and heatmap (**G**) of newly synthesized proteins identified in the H◊H, H◊L, and L→H conditions. **F-H**) Solar plots of H/L ratios for the three conditions H→H (**F**), H→L (**G**), L→H (**H**), highlighting major cellular organelles.

**SI Figure 3.**
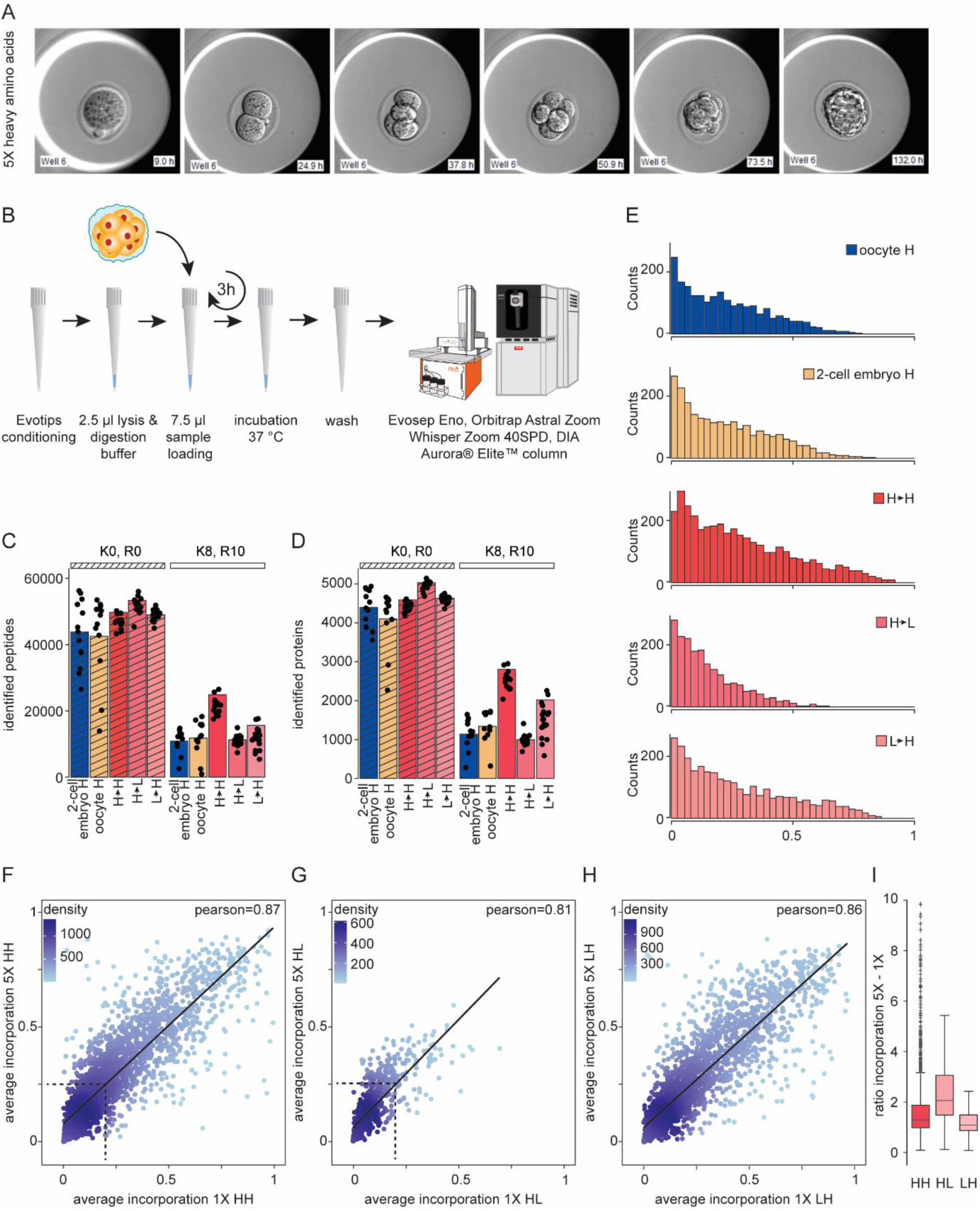
High correlation between the incorporation of heavy amino acids at 1X and 5X concentrations. **A**) Representative EmbryoScope images showing the development of zygotes over the first 130 h in medium containing 5X heavy amino acids. **B**) Schematic workflow, from left to right: Morulae were loaded onto preconditioned Evotips and spun down into the 3.5 µl of lysis and digestion buffer. After washing, samples were analyzed by an Evosep Eno coupled to an Orbitrap Astral Zoom LC-MS/MS setup. **C, D**) Numbers of identified peptides (**C**) and proteins (**D**) in the light (striped) and heavy channels. **E**) Histogram of incorporation values for each condition under the 5X heavy amino acid concentration. **F-H**) Density scatter plots comparing the incorporation of heavy amino acids between the 1X and 5X concentrations for the H◊H, H◊L, and L→H conditions. **I**) Bar plot showing a 2-2.5-fold higher incorporation at the 5X heavy amino acid concentration compared to 1X. H, heavy; L, light; EmbryoScope, time-lapse embryo imaging system.

**SI Figure 4.**
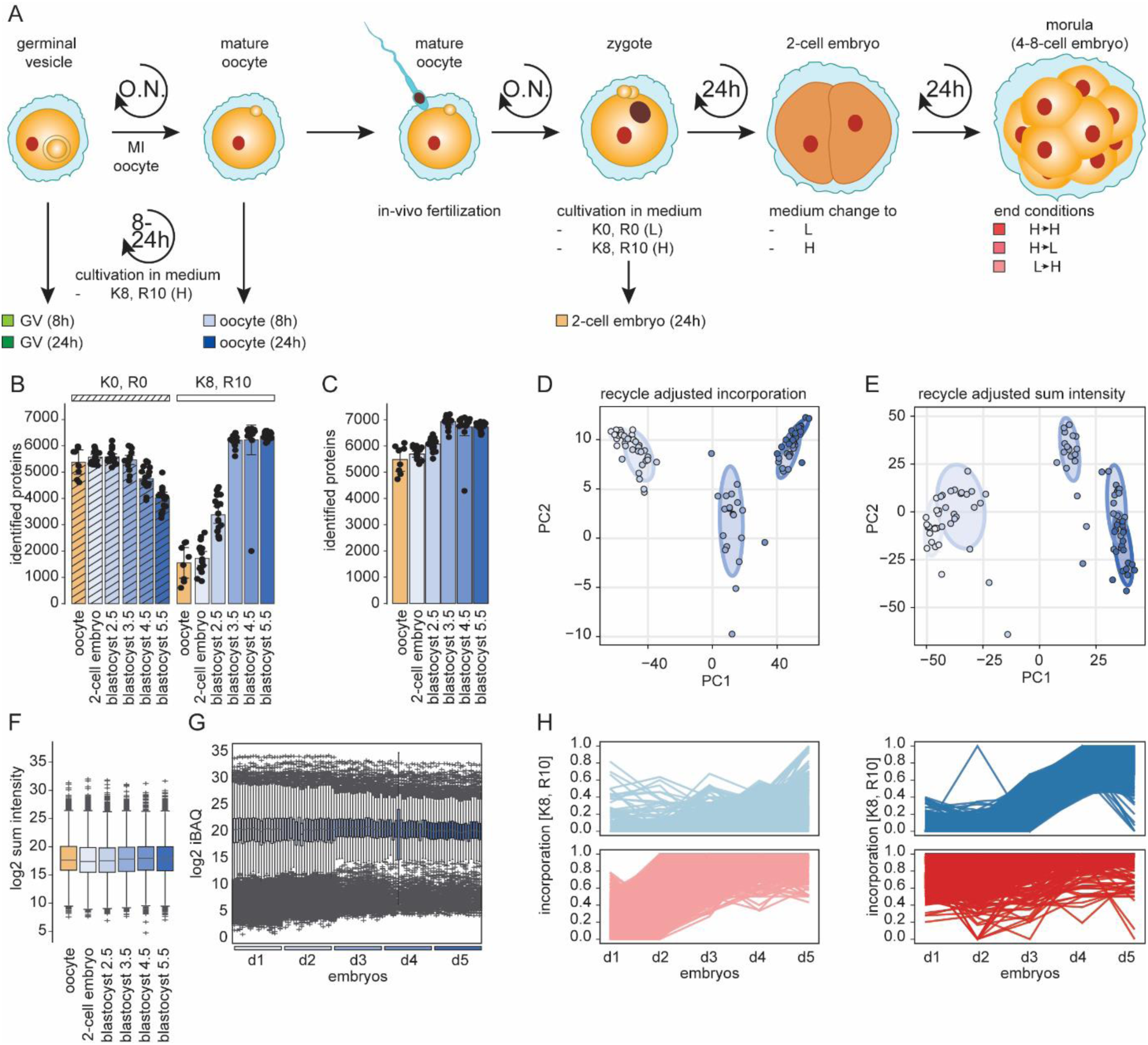
Heavy amino acid incorporation over five days. **A**) Schematic overview of the workflow for the cell culture experiment. From left to right: germinal vesicles, mature oocytes, or zygotes were isolated and cultured in either light [K0, R0] or heavy [K8, R10] SILAC medium. After 8, 24 h of incubation cells were harvested. **B**, **C**) Representative EmbryoScope pictures from the development of the first 130 h of zygotes in control medium (**B**) and in medium containing heavy amino acids (**C**). **D**) After washing, samples were analyzed by an Evosep One coupled to an Orbitrap Astral LC-MS/MS setup. **E**) Number of proteins identified in the light and heavy SILAC channels using DIA-NN for data analysis. **F**-**H**) Top identified GOterms for the incorporation, seperated to quartiles and for the three different conditions of H → H (**F**), L → H (**G**), and H → L (**H**). CTRL, control; DIA, data independent acquisition; H, heavy; LC-MS/MS, liquid chromatography tandem mass spectrometery; L, light; SILAC, stable isotope labeling in cell culture; O.N., overnight; SPD, samples per day.

**SI Figure 5.**
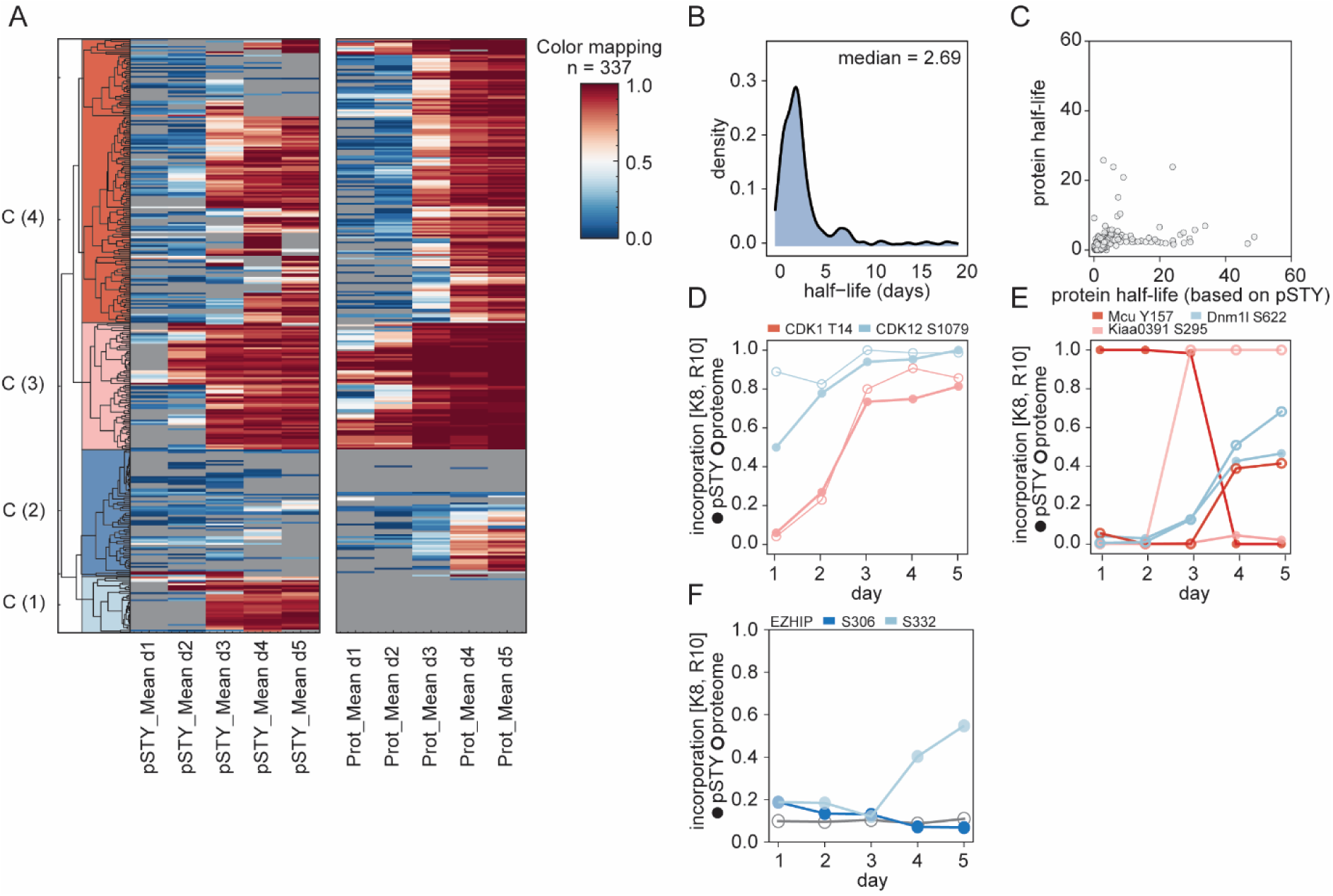
Phosphoproteome is following the trend of the proteome. **A)** Heatmap of the mean heavy amino acid incorporation of phospopeptides and proteins over 5 days. **B)** Density plot of the calculated half-life in days for proteins using the phosphopeptides. **C)** Scatter plot of the calculated protein half-life using the phophopeptides or proteins. **D-F**) Profile plots showing the incorporation of selected phosphosites and the respective protein.

**SI Figure 6.**
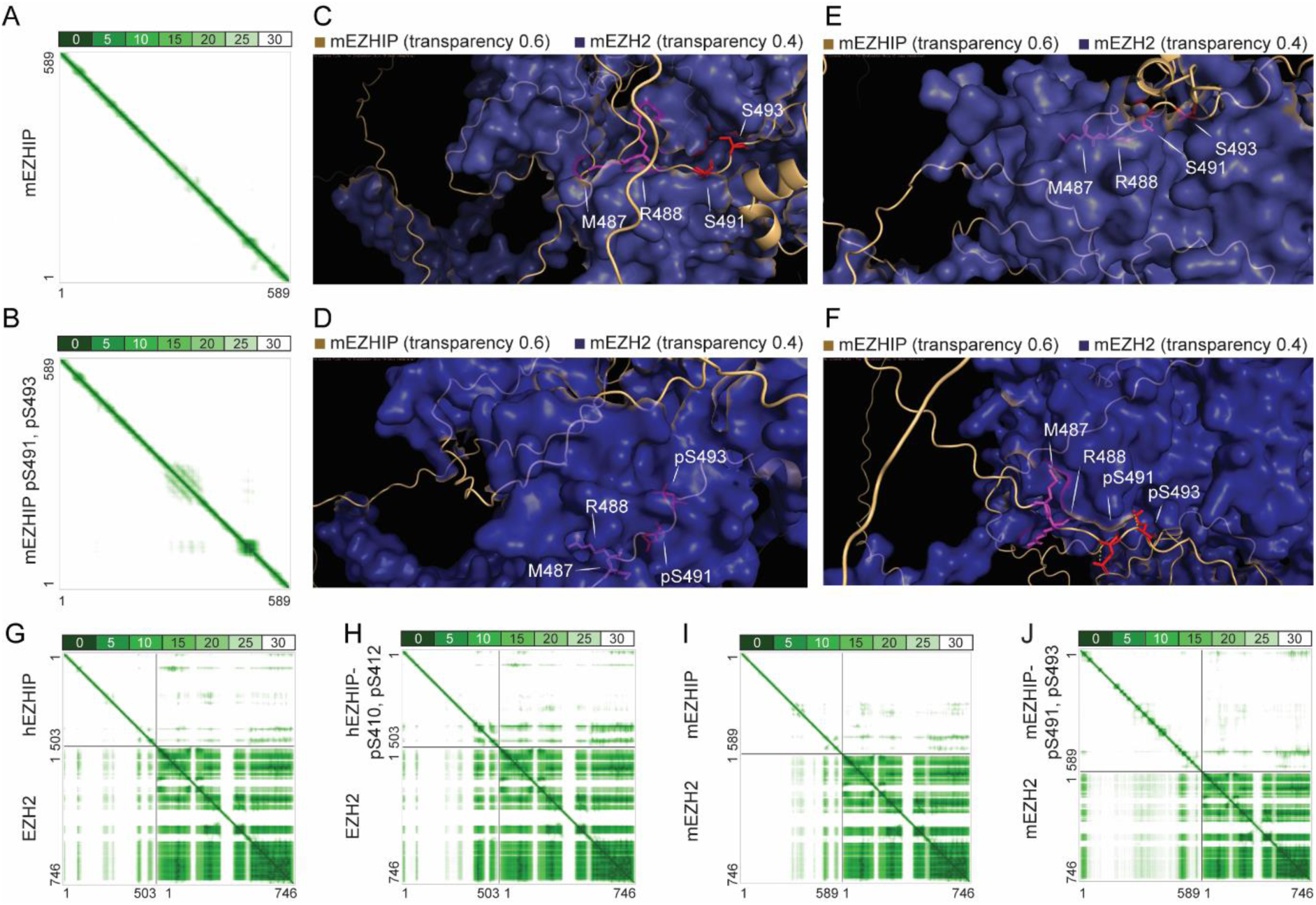
Phosphosites on the protein EZHIP changes the predicted structure of the interface between EYHIP and EZH2 in mice. A, **B**) Amino acid sequence matrix visualizing the EAD of mEZHIP without (**A**) and with (**B**) the phosphosites in the KLP motif. Structure and EAD prediction are based on AlphaFold. **C-F**) Zoomed in AlphaFold structures of mEZHIP-mEZH2 without (**C, E**) and with (**D, F**) the phosphosites in the KLP motif. M406 and R407, and (p)S410 and (p)S412 are visualized in purple and red stick form, respectively. Predicted polar contacts are visualized in a yellow, dashed line. EZH2 and EZHIP are visualized in purple surface and orange carton style, respectively. **G-J**) Amino acid sequence matrix visualizing the EAD of hEZHIP-hEZH2 and mEZHIP-mEZH2 without (**G, I**) and with (**H, J**) the phosphosites in the KLP motif. Structure and EAD prediction are based on AlphaFold.

**SI Figure 7.**
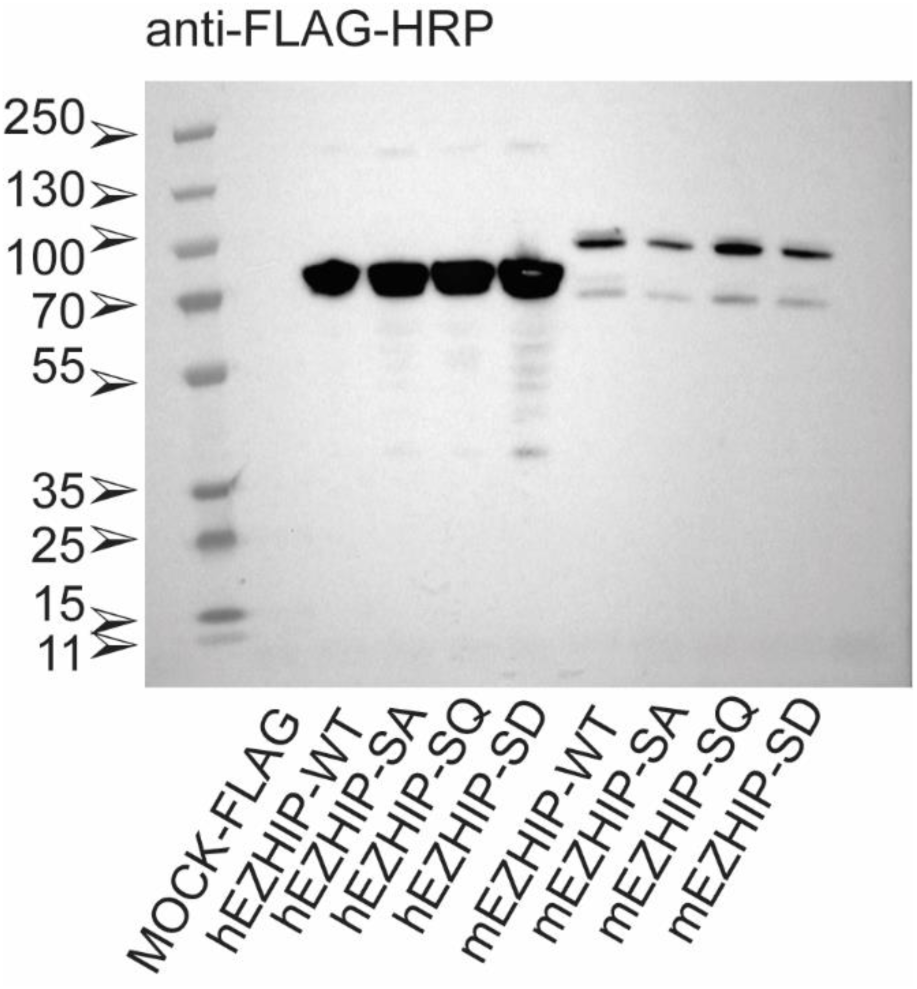
Expression of human and mouse EZHIP-FLAG constructs. Western blot analysis of transfected HEK cells with both human and mouse EZHIP-FLAG constructs.

## Declarations Ethics approval

This study complies with all relevant ethical regulations. Animal work was conducted according to license no. 2021-15-0201-00851, approved by the Danish National Animal Experiments Inspectorate, and performed according to national and local guidelines.

## Availability of data and materials

Code and EmbryoScope videos are available via Zenodo (https://zenodo.org/) the DOIs 10.5281/zenodo.22878819 and 10.5281/zenodo.22876670.

## Authors’ contributions

L.M.S., J.M.-G., and J.V.O. designed the experiments. L.M.S and M.L. prepared all samples and performed proteomics experiments. L.M.S., I.M., K.S., M.P.L., and M.L. prepared all cell culture work, L.M.S. analysed the resulting data. J.M.-G and J.H. performed animal handling and dissection, J.H., and M.L. performed ChiPseq data analysis, L.M.S. and I.A.H. performed MS method evaluation, L.M.S. wrote the first draft of the manuscript. E.H. and J.V.O. critically evaluated the results. All authors read, edited, and approved the final version of the manuscript.

## References

1. Aoki, F., Zygotic gene activation in mice: profile and regulation. J Reprod Dev, 2022. 68(2): p. 79–84.

2. Aoki, F., D.M. Worrad, and R.M. Schultz, Regulation of transcriptional activity during the first and second cell cycles in the preimplantation mouse embryo. Dev Biol, 1997. 181(2): p. 296–307.

3. Wassarman, P.M. and R.A. Kinloch, Gene expression during oogenesis in mice. Mutat Res, 1992. 296(1-2): p. 3–15.

4. Schultz, R.M., Regulation of zygotic gene activation in the mouse. Bioessays, 1993. 15(8): p. 531–8.

5. Zhao, J., et al., Dynamic metabolism during early mammalian embryogenesis. Development, 2023. 150(20).

6. Kojima, M.L., C. Hoppe, and A.J. Giraldez, The maternal-to-zygotic transition: reprogramming of the cytoplasm and nucleus. Nat Rev Genet, 2025. 26(4): p. 245–267.

7. Proks, M., N. Salehin, and J.M. Brickman, Deep learning-based models for preimplantation mouse and human embryos based on single-cell RNA sequencing. Nature Methods, 2025. 22(1): p. 207–216.

8. Yan, L., et al., Single-cell RNA-Seq profiling of human preimplantation embryos and embryonic stem cells. Nature Structural & Molecular Biology, 2013. 20(9): p. 1131–1139.

9. Zhao, C., et al., A comprehensive human embryo reference tool using single-cell RNA-sequencing data. Nature Methods, 2025. 22(1): p. 193–206.

10. Liu, Y., A. Beyer, and R. Aebersold, On the Dependency of Cellular Protein Levels on mRNA Abundance. Cell, 2016. 165(3): p. 535–50.

11. Upadhya, S.R. and C.J. Ryan, Experimental reproducibility limits the correlation between mRNA and protein abundances in tumor proteomic profiles. Cell Rep Methods, 2022. 2(9): p. 100288.

12. Fan, J., et al., Single-Cell RNA Sequencing Reveals Differences in Chromatin Remodeling and Energy Metabolism among In Vivo-Developed, In Vitro-Fertilized, and Parthenogenetically Activated Embryos from the Oocyte to 8-Cell Stages in Pigs. Animals, 2024. 14(3): p. 465.

13. Latham, K.E., Preimplantation embryo gene expression: 56 years of discovery, and counting. Mol Reprod Dev, 2023. 90(4): p. 169–200.

14. Vassena, R., et al., Waves of early transcriptional activation and pluripotency program initiation during human preimplantation development. Development, 2011. 138(17): p. 3699–709.

15. Domon, B. and R. Aebersold, Options and considerations when selecting a quantitative proteomics strategy. Nature Biotechnology, 2010. 28(7): p. 710–721.

16. Mann, M., Fifteen years of Stable Isotope Labeling by Amino Acids in Cell Culture (SILAC). Methods Mol Biol, 2014. 1188: p. 1–7.

17. Ong, S.-E., et al., Stable Isotope Labeling by Amino Acids in Cell Culture, SILAC, as a Simple and Accurate Approach to Expression Proteomics*. Molecular & Cellular Proteomics, 2002. 1(5): p. 376–386.

18. Ong, S.E. and M. Mann, A practical recipe for stable isotope labeling by amino acids in cell culture (SILAC). Nat Protoc, 2006. 1(6): p. 2650–60.

19. Cargile, B.J., et al., Synthesis/degradation ratio mass spectrometry for measuring relative dynamic protein turnover. Anal Chem, 2004. 76(1): p. 86–97.

20. He, Y., et al., On-capillary alkylation micro-reactor: a facile strategy for proteo-metabolome profiling in the same single cells. Chem Sci, 2023. 14(46): p. 13495–13502.

21. Ye, Z., et al., One-Tip enables comprehensive proteome coverage in minimal cells and single zygotes. Nature Communications, 2024. 15(1): p. 2474.

22. Zhang, H., et al., Deep Profiling of Oocyte Aging Enabled by Simple One-Step Vial-Based Pretreatment and Single-Cell Proteomics. JACS Au, 2025. 5(5): p. 2321–2333.

23. Bubis, J.A., et al., Challenging the Astral mass analyzer to quantify up to 5,300 proteins per single cell at unseen accuracy to uncover cellular heterogeneity. Nat Methods, 2025. 22(3): p. 510–519.

24. Guzman, U.H., et al., Ultra-fast label-free quantification and comprehensive proteome coverage with narrow-window data-independent acquisition. Nature Biotechnology, 2024. 42(12): p. 1855–1866.

25. Hendriks, I.A., et al., Extending single-cell proteomics from HeLa to small primary human immune cells. Nature Communications, 2026.

26. Demichev, V., et al., DIA-NN: neural networks and interference correction enable deep proteome coverage in high throughput. Nature Methods, 2020. 17(1): p. 41–44.

27. Derks, J., et al., Increasing the throughput of sensitive proteomics by plexDIA. Nature Biotechnology, 2023. 41(1): p. 50–59.

28. Sabatier, P., et al., Global analysis of protein turnover dynamics in single cells. Cell, 2025. 188(9): p. 2433–2450.e21.

29. Ozadam, H., et al., Single-cell quantification of ribosome occupancy in early mouse development. Nature, 2023. 618(7967): p. 1057–1064.

30. Santon, J.B. and M. Pellegrini, Rates of ribosomal protein and total protein synthesis during Drosophila early embryogenesis. Developmental Biology, 1981. 85(1): p. 252–257.

31. Hsu, C., et al., Evaluation of a prototype Orbitrap Astral Zoom mass spectrometer for quantitative proteomics - Beyond identification lists. bioRxiv, 2025.

32. Guzman, U.H., et al., Higher-Throughput Proteome Profiling Enabled by Parallelized Pre-Accumulation and Optimized Ion Processing in the Orbitrap Astral Zoom Mass Spectrometer. bioRxiv, 2025: p. 2025.07.14.664789.

33. Li, L., B. Baibakov, and J. Dean, A Subcortical Maternal Complex Essential for Preimplantation Mouse Embryogenesis. Developmental Cell, 2008. 15(3): p. 416–425.

34. Bebbere, D., et al., The subcortical maternal complex: emerging roles and novel perspectives. Mol Hum Reprod, 2021. 27(7).

35. Bebbere, D., et al., Expression of maternally derived KHDC3, NLRP5, OOEP and TLE6is associated with oocyte developmental competence in the ovine species. BMC Developmental Biology, 2014. 14(1): p. 40.

36. Jentoft, I.M.A., et al., Mammalian oocytes store proteins for the early embryo on cytoplasmic lattices. Cell, 2023. 186(24): p. 5308–5327.e25.

37. Leese, H.J., P.J. McKeegan, and R.G. Sturmey, Amino Acids and the Early Mammalian Embryo: Origin, Fate, Function and Life-Long Legacy. Int J Environ Res Public Health, 2021. 18(18).

38. Beynon, R.J. and J.M. Pratt, Metabolic labeling of proteins for proteomics. Mol Cell Proteomics, 2005. 4(7): p. 857–72.

39. Leduc, A., et al., *Principles* of protein abundance regulation across single cells in a mammalian tissue. bioRxiv, 2025.

40. Schwanhäusser, B., et al., Global quantification of mammalian gene expression control. Nature, 2011. 473(7347): p. 337–42.

41. Wu, C.C., et al., Metabolic labeling of mammalian organisms with stable isotopes for quantitative proteomic analysis. Anal Chem, 2004. 76(17): p. 4951–9.

42. Christen, P., R. Jaussi, and R. Benoit, Stoffwechsel der Proteine und Aminosäuren, in Biochemie und Molekularbiologie: Eine Einführung in 40 Lerneinheiten, P. Christen, R. Jaussi, and R. Benoit, Editors. 2024, Springer Berlin Heidelberg: Berlin, Heidelberg. p. 279–302.

43. Hornbeck, P.V., et al., PhosphoSitePlus, 2014: mutations, PTMs and recalibrations. Nucleic Acids Res, 2015. 43(Database issue): p. D512–20.

44. Chen, M., et al., EPSD 2.0: An Updated Database of Protein Phosphorylation Sites Across Eukaryotic Species. Genomics Proteomics Bioinformatics, 2025. 23(3).

45. Raes, A., et al., Cathepsin-L Secreted by High-Quality Bovine Embryos Exerts an Embryotrophic Effect In Vitro. Int J Mol Sci, 2023. 24(7).

46. ZHU Qing, S.J., TANG Huai-yun, The Role of Cathepsin L in Oocyte and Early Embryonic Development. Journal of International Reproductive Health/Family Planning, 2026. 45(1): p. 49–53.

47. Hayakawa, K., et al., Oocyte-specific linker histone H1foo is an epigenomic modulator that decondenses chromatin and impairs pluripotency. Epigenetics, 2012. 7(9): p. 1029–36.

48. Szklarczyk, D., et al., The STRING database in 2025: protein networks with directionality of regulation. Nucleic Acids Res, 2025. 53(D1): p. D730–d737.

49. Satouh, Y. and K. Sato, Reorganization, specialization, and degradation of oocyte maternal components for early development. Reproductive Medicine and Biology, 2023. 22(1): p. e12505.

50. Tsukamoto, S. and T. Tatsumi, Degradation of maternal factors during preimplantation embryonic development. J Reprod Dev, 2018. 64(3): p. 217–222.

51. Hu, Z., et al., CKAP5 deficiency induces premature ovarian insufficiency. EBioMedicine, 2025. 115: p. 105718.

52. Zhang, N., T. Wakai, and R.A. Fissore, Caffeine alleviates the deterioration of Ca(2+) release mechanisms and fragmentation of in vitro-aged mouse eggs. Mol Reprod Dev, 2011. 78(9): p. 684–701.

53. Khan, I., et al., Time of insemination and its effect on in-vitro fertilization, cleavage and pregnancy rates in GnRH agonist/HMG-stimulated cycles. Hum Reprod, 1989. 4(8): p. 921–6.

54. Igarashi, H., et al., Aging-related changes in calcium oscillations in fertilized mouse oocytes. Mol Reprod Dev, 1997. 48(3): p. 383–90.

55. Chen, J., et al., Exposure to Copper Compromises the Maturational Competency of Porcine Oocytes by Impairing Mitochondrial Function. Front Cell Dev Biol, 2021. 9: p. 678665.

56. Jain, S.U., et al., PFA ependymoma-associated protein EZHIP inhibits PRC2 activity through a H3 K27M-like mechanism. Nature Communications, 2019. 10(1): p. 2146.

57. Ragazzini, R., et al., EZHIP constrains Polycomb Repressive Complex 2 activity in germ cells. Nature Communications, 2019. 10(1): p. 3858.

58. Kang, J., et al., Polycomb Repressive-Deubiquitinase Complex Safeguards Oocyte Epigenome and Female Fertility by Restraining Polycomb Activity. bioRxiv, 2025.

59. Miller, S.A., et al., Full methylation of H3K27 by PRC2 is dispensable for initial embryoid body formation but required to maintain differentiated cell identity. Development, 2021. 148(7).

60. Liu, N. and B. Zhu, Chapter 10 - Regulation of PRC2 Activity, in Polycomb Group Proteins, V. Pirrotta, Editor. 2017, Academic Press. p. 225–258.

61. Hübner, J.M., et al., EZHIP/CXorf67 mimics K27M mutated oncohistones and functions as an intrinsic inhibitor of PRC2 function in aggressive posterior fossa ependymoma. Neuro Oncol, 2019. 21(7): p. 878–889.

62. Jumper, J., et al., Highly accurate protein structure prediction with AlphaFold. Nature, 2021. 596(7873): p. 583–589.

63. Halliwell, J.A., et al., Sex-specific DNA-replication in the early mammalian embryo. Nature Communications, 2024. 15(1): p. 6323.

64. Liu, X., et al., Distinct features of H3K4me3 and H3K27me3 chromatin domains in pre-implantation embryos. Nature, 2016. 537(7621): p. 558–562.

65. Diop, S., et al., H3K27me3-dependent imprinting and transcriptional regulation in early mouse embryos requires EZHIP-mediated restriction of PRC2 activity. Nat Commun, 2026. 17(1): p. 1758.

66. Zeng, Y., et al., EZHIP restricts noncanonical PRC2 binding and regulates H3K27me3 intergenerational inheritance and reprogramming. Cell Stem Cell, 2025. 32(11): p. 1741–1757.e5.

67. Ajuyah, P., et al., Histone H3-wild type diffuse midline gliomas with H3K27me3 loss are a distinct entity with exclusive EGFR or ACVR1 mutation and differential methylation of homeobox genes. Sci Rep, 2023. 13(1): p. 3775.

68. Batth, T.S., et al., Protein Aggregation Capture on Microparticles Enables Multipurpose Proteomics Sample Preparation. Mol Cell Proteomics, 2019. 18(5): p. 1027–1035.

69. Bekker-Jensen, D.B., et al., A Compact Quadrupole-Orbitrap Mass Spectrometer with FAIMS Interface Improves Proteome Coverage in Short LC Gradients. Mol Cell Proteomics, 2020. 19(4): p. 716–729.

70. Martinez-Val, A., et al., Spatial-proteomics reveals phospho-signaling dynamics at subcellular resolution. Nature Communications, 2021. 12(1): p. 7113.

71. Griffith, K.A., et al., African Americans with a family history of colorectal cancer: barriers and facilitators to screening. Oncol Nurs Forum, 2012. 39(3): p. 299–306.

72. Chen, S., et al., *fastp: an ultra-fast all-in-one FASTQ preprocessor*. Bioinformatics, 2018. 34(17): p. i884–i890.

73. Langmead, B. and S.L. Salzberg, Fast gapped-read alignment with Bowtie 2. Nat Methods, 2012. 9(4): p. 357–9.

74. Li, H., et al., The Sequence Alignment/Map format and SAMtools. Bioinformatics, 2009. 25(16): p. 2078–9.

75. Ewels, P., et al., MultiQC: summarize analysis results for multiple tools and samples in a single report. Bioinformatics, 2016. 32(19): p. 3047–8.

76. Lee, C.C., W.C. Su, and W.C. Chang, Effects of the charge-transfer reorganization energy on the open-circuit voltage in small-molecular bilayer organic photovoltaic devices: comparison of the influence of deposition rates of the donor. Phys Chem Chem Phys, 2016. 18(18): p. 12651–61.

